# One-Hot News: Drug Synergy Models Shortcut Molecular Features

**DOI:** 10.1101/2025.04.18.649584

**Authors:** Emine Beyza Çandır, Halil İbrahim Kuru, Magnus Rattray, A. Ercument Cicek, Oznur Tastan

## Abstract

Combinatorial drug therapy holds great promise for tackling complex diseases, but the vast number of possible drug combinations makes exhaustive experimental testing infeasible. Computational models have been developed to guide experimental screens by assigning synergy scores to drug pair–cell line combinations, where they take input structural and chemical information on drugs and molecular features of cell lines. The premise of these models is that they leverage this biological and chemical information to predict synergy measurements. In this study, we demonstrate that replacing drug and cell line representations with simple one-hot encodings results in comparable or even slightly improved performance across diverse published drug combination models. This unexpected finding suggests that current models use these representations primarily as identifiers and exploit covariation in the synergy labels. Our synthetic data experiments show that models can learn from the true features; however, when drugs and cell lines recur across drug–drug–cell triplets, this repeating structure impairs feature-based learning. While the current synergy prediction models can still aid in prioritizing drug pairs within a panel of tested drugs and cell lines, our results highlight the need for better strategies to learn from intended features and generalize to unseen drugs and cell lines.

## Introduction

Synergistic drug combinations enable lower dosing of each agent, reducing adverse side effects, which is especially important in treating complex diseases such as cancer (Mokhtari *et al*., 2017; Al-Lazikani *et al*., 2012; Tamma *et al*., 2012; Möttönen *et al*., 1999; Gradman et al., 2010). However, the vast number of possible drug pairs and cell line combinations makes exhaustive clinical evaluations of drug combinations infeasible. To address this limitation, several computational models have been developed to guide experimental efforts (Abbasi and Rousu, 2024). By predicting synergistic scores for drug pairs on cell lines, these models help prioritize which combinations merit further testing.

In a typical synergy prediction model, each input example is composed of a triplet: a drug pair and the cell line on which the two drugs’ combined effect is measured. Models take numerical representations of these triplets using various descriptors for drugs and cell lines. For example, to describe drugs, many methods employ chemical and structural fingerprints (Xu *et al*., 2023; Wang *et al*., 2023b; Preuer *et al*., 2017; Kuru *et al*., 2022; Güvenç Paltun et al., 2021). Graph-based representations have also been used, such as in DeepDDS (Wang *et al*., 2021). There are also multimodal approaches that use different representations jointly, such as JointSyn (Li *et al*., 2024), which uses fingerprints and molecular graphs. Alternative strategies have been proposed; for example, Marsy captures drug characteristics through differential expression signatures induced in two specific cell lines (El Khili *et al*., 2023). The biological context of cell lines is often represented by the untreated gene expression profiles of the cell line. The premise of all these models is that they intend to leverage the chemical and biological information of drugs and cell lines to predict synergy.

Previously, models have been reported to perform well on new combinations of drugs and cell lines that are exclusively seen during training but fail on new drugs or cell lines during testing. This is revealed in the performances obtained in evaluation setups where the test data are split in a stratified way (Li *et al*., 2024; Abbasi and Rousu, 2024; Wang *et al*., 2024). These split scenarios for drug-drug-cell line combinations are illustrated in Figure 1 and are detailed in the Methods. Models perform very well on unseen triplet examples (leave-triplet-out) as this setup is relatively easy. High performance is also reported for the unseen drug pairs (leave-drug-pair-out), where both drugs are observed in training but only within triplets where they are paired with other drugs and cell lines.

**Fig. 1.**
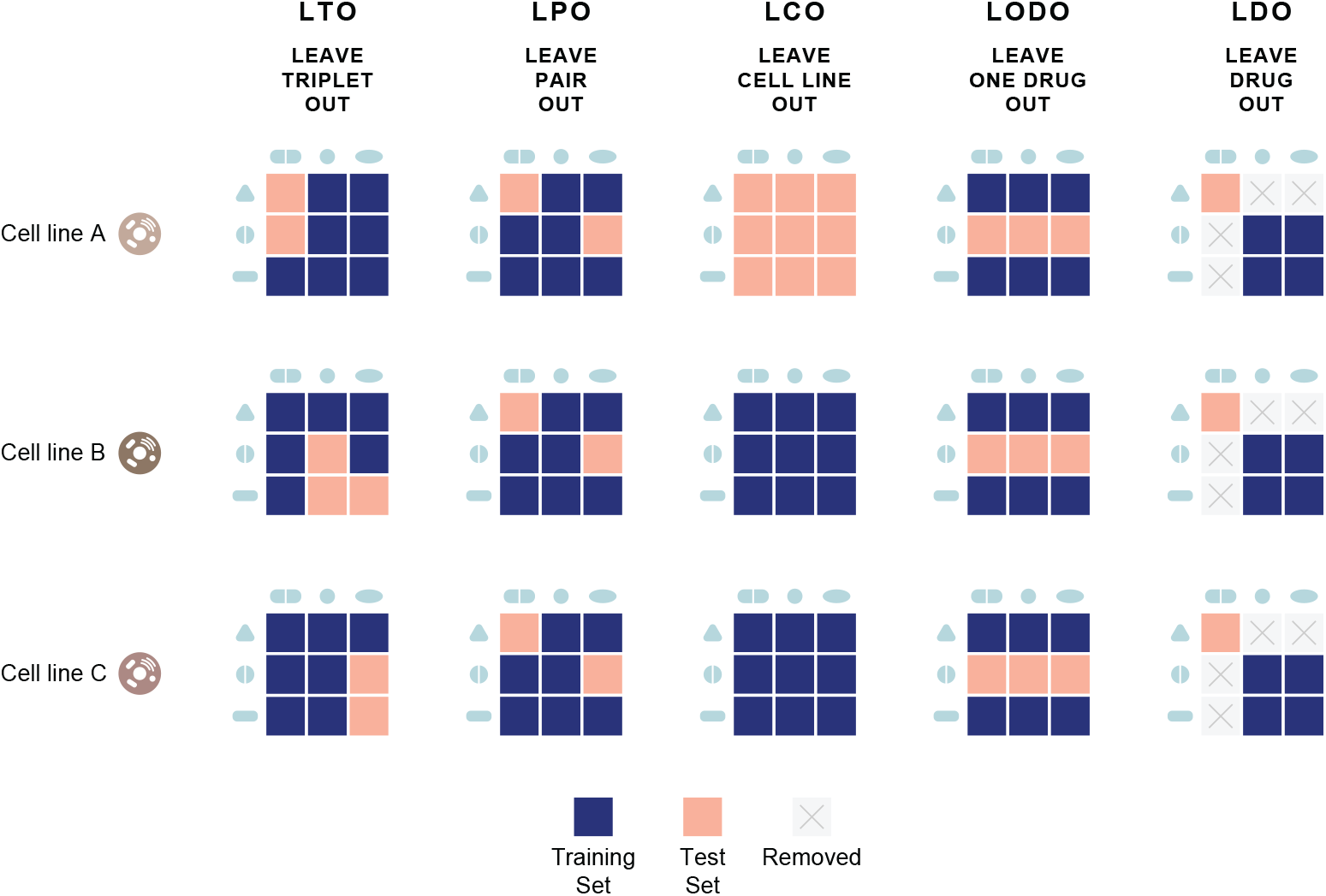
Illustration of data split strategies, inspired by Preuer *et al*. (2017). Each evaluation strategy shows how drugs and cell lines are included or excluded in the training and test sets. The detailed description of these strategies is explained in Section 2.4.

Model performance deteriorates when models are evaluated in triplets of an unseen cell line (leave-cell-line-out) (Preuer *et al*., 2017; Xu *et al*., 2023; Li *et al*., 2024; Zhang and Tu, 2023). Model predictive performances also decrease when the test dataset is formed of triplets where one of the tested drugs is not observed in the training (leave-one-drug-out) (Preuer *et al*., 2017; Xu et al., 2023; Zhang and Tu, 2023). In particular, all model performances suffer when they are evaluated on triplets where neither of the drugs in the triplet is observed during training (Li *et al*., 2024).

While a generalization problem has been reported in the literature, we provide direct evidence that many drug–synergy models may not be using the intended chemical and biological information. Across a diverse set of published architectures, we replaced the original drug and cell-line representations with simple one-hot encodings—and performance stayed largely unchanged, and in some cases even improved. Because one-hot vectors are merely unique, orthogonal IDs with no biological content, this result suggests that models often rely on identity-linked covariation rather than biochemical features. We verified this by reusing the authors’ public code and their original data splits for each method, observing the same pattern consistently. Our goal is not to rank architectures or encodings, but to test whether each model actually uses its designated inputs. To that end, we held splits, hyperparameters, and training protocols fixed and swapped the biological descriptors for one-hot vectors, thereby directly measuring how much performance depends on biologically informed representations under tightly controlled conditions.

While previous work has documented generalization issues when the models are applied to unseen drugs and cell lines, our results reveal a more fundamental issue than previously acknowledged. The models simply learn from the covariation patterns of the synergy measurements, shortcutting the biologically relevant features, which could be the reason behind the generalization barrier. Through synthetic data experiments, we investigate under what conditions models can base their predictions on the true features. We find that the repeat structure of the data impairs the feature-based learning. Below, we describe in detail the experiments we conducted.

## Methods

### Tested Models

In our analysis, we selected different models that are well-recognized or recent: DeepSynergy (Preuer *et al*., 2017), DeepDDS (Wang *et al*., 2021), MatchMaker (Kuru *et al*., 2022), MARSY (El Khili *et al*., 2023), and JointSyn (Li *et al*., 2024). Each model uses different approaches and features for drug synergy prediction, while all use cell line gene expression profiles to represent the biological context. Below, we provide a summary of these approaches. The details about the number of features and the dataset descriptions are also provided in Table 1.

**Table 1.**
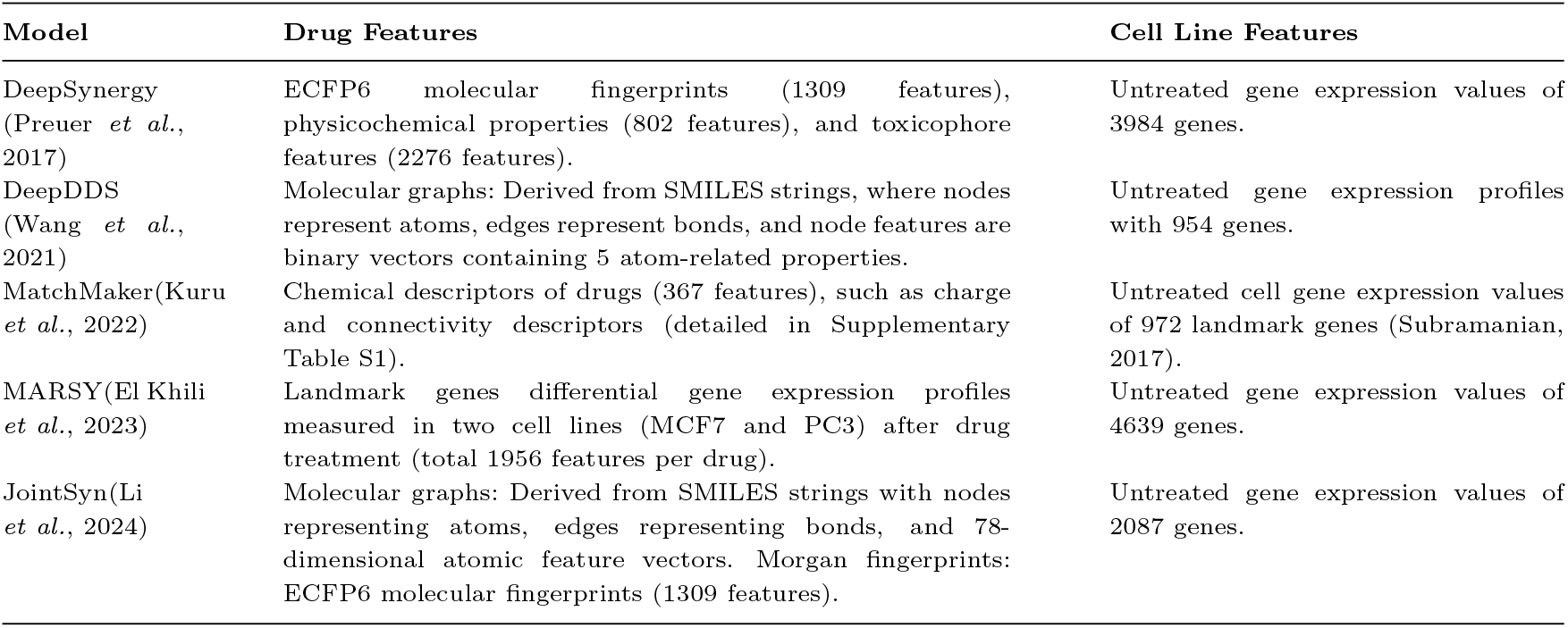
Summary of Drug and Cell Line Features Used by Each Model.

| Model | Drug Features | Cell Line Features |
| --- | --- | --- |
| DeepSynergy (Preuer *et al.*, 2017) | ECFP6 molecular fingerprints (1309 features), physicochemical properties (802 features), and toxicophore features (2276 features). | Untreated gene expression values of 3984 genes. |
| DeepDDS (Wang *et al.*, 2021) | Molecular graphs: Derived from SMILES strings, where nodes represent atoms, edges represent bonds, and node features are binary vectors containing 5 atom-related properties. | Untreated gene expression profiles with 954 genes. |
| MatchMaker(Kuru *et al.*, 2022) | Chemical descriptors of drugs (367 features), such as charge and connectivity descriptors (detailed in Supplementary Table S1). | Untreated cell gene expression values of 972 landmark genes (Subramanian, 2017). |
| MARSY(El Khili *et al.*, 2023) | Landmark genes differential gene expression profiles measured in two cell lines (MCF7 and PC3) after drug treatment (total 1956 features per drug). | Untreated gene expression values of 4639 genes. |
| JointSyn(Li *et al.*, 2024) | Molecular graphs: Derived from SMILES strings with nodes representing atoms, edges representing bonds, and 78-dimensional atomic feature vectors. Morgan fingerprints: ECFP6 molecular fingerprints (1309 features). | Untreated gene expression values of 2087 genes. |

- **DeepSynergy** pioneered the use of deep learning for synergy prediction (Preuer *et al*., 2017). It uses a fully connected neural network that concatenates feature vectors for two drugs and a cell line. Each drug is represented by three types of chemical descriptors: extended connectivity fingerprints (ECFP6) with 1309 features, physicochemical properties with 802 features, and 2276 toxicophore features. To characterize cell lines, DeepSynergy uses gene expression profiles of untreated cells (Iorio, 2016), filtered down to 3984 informative genes.
- **DeepDDS** is a GNN-based model designed to learn drug representations from molecular graphs created from SMILES strings. In these graphs, nodes represent atoms, and edges represent bonds. Each node is described by a binary vector containing five atomic properties. For cell lines, DeepDDS uses baseline gene expression profiles of untreated cells, filtered to include 954 genes. These genes were selected by intersecting CCLE expression data with LINCS landmark genes and removing noncoding RNA transcripts (Wang *et al*., 2021).
- **MatchMaker** employs two parallel drug-specific subnetworks alongside a Synergy Prediction Network (SPN), where each drug-specific subnetwork (DSN) processes the chemical features of one drug and the gene expression features of the corresponding cell line (Kuru *et al*., 2022). The outputs of the DSNs are concatenated and fed into the SPN to predict synergy scores. Drugs are represented using 367 chemical descriptors calculated with the PyBioMed library. These descriptors encode the chemical structure of each drug. For cell lines, baseline gene expression values of untreated cells, consisting of 972 landmark genes, are used.
- **MARSY** generates representations for drug pairs and their interactions with cell lines through separate encoders (El Khili *et al*., 2023). These representations are integrated within a multitask predictor to output both synergy scores and single-drug responses. MARSY represents drugs using their differential gene expression (DGE) signatures obtained from the LINCS database (Subramanian, 2017). These signatures were measured in two cell lines, MCF7 and PC3, 24 hours after drug treatment. Each drug was characterized by a concatenation of 978 landmark genes from both cell lines, resulting in 1,956 features per drug. For cell lines, MARSY utilizes baseline gene expression profiles of untreated cells from the CCLE via the CellMiner database (Reinhold *et al*., 2012). After filtering out lowly expressed and low-variance genes, the final cell line representation includes 4,639 genes.
- **JointSyn** is a recent model that reported competitive performance compared to other state-of-the-art methods (Li *et al*., 2024). It integrates a joint graph encompassing drug combinations, drug features, and cell line representations within a dual-view architecture. These views generate embeddings for both the drug combination and the cell line, which are then passed to a prediction network to estimate synergy scores. JointSyn represents drugs using molecular graphs and Morgan fingerprints. Molecular graphs are derived from SMILES strings using RDKit (Landrum, 2016), where atoms are represented as nodes and bonds as edges. Each node is characterized by a 78-dimensional atomic feature vector computed with DeepChem (Ramsundar *et al*., 2019). Additionally, Morgan fingerprints with 1,309 features capture further structural characteristics. For cell lines, JointSyn employs baseline gene expression profiles comprising 2,087 genes, which are filtered from the CCLE database based on drug sensitivity relevance.

### Datasets and processing

Our objective is not to benchmark models on a uniform dataset and determine the best performing one, but to test *within each model* whether performance depends on its intended biological features. To this end, we deliberately keep each method’s original published pipeline (splits, hyperparameters, training) fixed (except for MatchMaker, which we explain below), and change only the input representation (original features vs. one-hot). Below, we provide the details of each model’s dataset. Synergy datasets comprise triplets: a drug pair and the cell lines, along with a synergy score derived from an experimental grid of measurements made at different drug dosages.

DeepSynergy was evaluated using the O’Neil dataset, available on the DeepSynergy website, which includes 23,052 drug–cell line combinations involving 38 drugs and 39 cell lines. Similarly, the DeepDDS model used another subset of the O’Neil dataset. This set included 12,415 combinations involving 36 drugs and 31 cell lines. JointSyn also uses a subset of the O’Neil dataset, as detailed in its publication, comprising 12,033 combinations of 38 drugs and 34 cell lines. MARSY was trained on a filtered version of the DrugComb dataset provided by its authors, containing 86,348 combinations involving 670 drugs and 75 cell lines.

For the MatchMaker model, we were able to conduct a more in-depth analysis. We used both the DrugComb and NCI-ALMANAC datasets. We employed an updated version of the DrugComb dataset, which was different from the original publication. We used this larger dataset because it is more suitable for the leave-drug-out split, wherein a subset of the data must be excluded. The DrugComb dataset comprises 739,964 drug-drug–cell line combinations involving 8,397 drugs and 2,320 cell lines. We filtered this dataset to include only drugs with available structural information in the PubChem database and cell lines with accessible gene expression data from the Genomics of Drug Sensitivity in Cancer (GDSC) database (Iorio, 2016). Following this filtering process, we obtained 426,239 combinations covering 3,057 drugs and 162 cell lines. The DrugComb dataset is imbalanced in the frequency with which individual drugs are represented across combinations. Approximately two-thirds of the drugs are involved in only one or two combinations within the dataset. In contrast, some drugs appear in more than ten thousand combinations. This imbalance extends to drug pairs as well; as illustrated in Supplementary Figure S1, one drug in a pair may be present in a single combination, while another one is included in over 7,000 combinations. Given the imbalanced nature of the DrugComb dataset, which may contribute to the results, we extended our experiments to include the NCI-ALMANAC dataset, which is more balanced. The original NCI-ALMANAC dataset comprised 304,549 combinations of 104 drugs and 60 cell lines. After applying the same filtering criteria, based on drug structures and cell line gene expressions we used for DrugComb, there were 264,424 combinations involving 99 drugs and 54 cell lines. As illustrated in Supplementary Figure S2, over 75% of drug pairs in the filtered dataset feature both drugs appearing between 5,000 and 6,000 times. Additionally, every drug in the dataset appears in at least 4,000 combinations, resulting in a more evenly distributed dataset.

The models are trained to predict different synergy scores. MatchMaker uses the Loewe Additivity score (Loewe, 1953) when DrugComb is employed. When trained on NCI-ALMANAC MatchMaker, the ComboScore is employed instead of Loewe. MARSY uses the ZIP score from the DrugComb dataset. DeepDDS applies a threshold on the Loewe Additivity score to binarize. Combinations with a synergy score greater than 10 are classified as synergistic, while scores below zero are categorized as antagonistic.

Table 2 summarizes these datasets used in our experiments for each model, including the synergy score metrics, as well as the number of drug combinations, drugs, and cell lines. For the one-hot encoded models, the union of drugs in the train and test is used to form the dictionary of drugs and the one-hot encoding vectors are created. The same is repeated for the cell lines.

**Table 2.** Summary of Synergy Scores and Dataset Statistics for Each Model. Note that while some models use datasets from the same sources, the sizes of these datasets vary according to the distinct filtering criteria employed in each study.

| Models - Dataset | Synergy Score | # of Combinations | # of Drugs | # of Cell Lines |
| --- | --- | --- | --- | --- |
| DeepSynergy - O’Neil | Loewe | 23,052 | 38 | 39 |
| DeepDDS - O’Neil | Binarized Loewe | 12,415 | 36 | 31 |
| MatchMaker - DrugComb | Loewe | 426,239 | 3,057 | 162 |
| MatchMaker - NCI-ALMANAC | ComboScore | 264,424 | 99 | 54 |
| MARSY - DrugComb | Zip | 86,348 | 670 | 75 |
| JointSyn - O’Neil | Loewe | 12,033 | 38 | 34 |

**Table 3.** Performance Comparison of MatchMaker Model Using Drug & Cell Line Features vs OHE Representations on DrugComb Dataset Across Different Split Methods. Chemical descriptors were originally used to represent drug features, while cell line gene expression levels represented cell line features in the MatchMaker model. These representations serve as the baseline for comparison with one-hot-encoded drug and cell line representations.

| Split Method | Feature Type | MSE | PCC | SCC |
| --- | --- | --- | --- | --- |
| LPO | Drug & Cell Feature | 97.23 ± 1.14 | 0.76 ± 0.002 | 0.72 ± 0.001 |
| | One-Hot-Encoding | 100.25 ± 1.15 | 0.75 ± 0.002 | 0.70 ± 0.002 |
| LCO | Drug & Cell Feature | 161.68 ± 1.50 | 0.52 ± 0.013 | 0.45 ± 0.012 |
| | One-Hot-Encoding | 158.11 ± 1.45 | 0.53 ± 0.012 | 0.46 ± 0.012 |
| LODO | Drug & Cell Feature | 199.22 ± 1.75 | 0.39 ± 0.009 | 0.36 ± 0.007 |
| | One-Hot-Encoding | 202.16 ± 1.76 | 0.33 ± 0.009 | 0.28 ± 0.007 |
| LDO | Drug & Cell Feature | 237.42 ± 3.34 | 0.13 ± 0.012 | 0.14 ± 0.010 |
| | One-Hot-Encoding | 223.57 ± 3.18 | 0.10 ± 0.009 | 0.10 ± 0.008 |

### Other Baselines and Compared Representations

In addition to the OHE baseline, we evaluated several complementary baselines and a model that leverages pretrained molecular representations.

#### Average baseline predictors

We implemented four simple, average-based regressors: (i) Overall Average—predict the global mean synergy across all training samples for every test instance; (ii) Drug-Pair Average—if the exact drug pair appears in training, predict its mean synergy; otherwise, use the overall average; (iii) Cell-Line Average—if the test cell line appears in training, predict its mean synergy; otherwise, use the overall average; (iv) Cell-Line & At-Least-One-Drug Average—if the test cell line and at least one of its drugs appear in training, predict the mean synergy for that combination; otherwise, use the overall average.

#### Shuffled features

As a control, we randomly permuted the entries of drug and cell-line feature vectors across samples, preserving feature distributions while breaking entity-specific associations.

#### MoLFormer (pretrained drug embeddings)

Drugs were encoded using pretrained MoLFormer-XL-both-10pct embeddings released by IBM Research and available via the HuggingFace Transformers library (Ross *et al*., 2022). Cell lines were represented with one-hot vectors. Drug embeddings were obtained by tokenizing the SMILES representation of each compound and passing it through the pretrained MoLFormer model. The pooled output vector from the model was used as the fixed-length embedding for each drug, resulting in a 768-dimensional representation.

### Data Splitting Strategies for Evaluations

The drug combination model performance can be assessed in different evaluation sets, where the criteria for the test dataset differ. These scenarios affect the prediction task difficulty and are important to understand the model’s capabilities and performance claims. We illustrate these scenarios in Figure 1.

- **Leave-Triple-Out (LTO):** The drug-drug–cell line triplets are randomly split without any stratification. The test set includes unseen drug-drug–cell line triplets. However, individual drugs, drug pairs or cell lines within these combinations may still appear with another partner in the training data. This is the easiest scenario.
- **Leave-Pair-Out (LPO):** Drug pairs present in the test set are not included in the training set. However, individual drugs within these pairs can still appear in the training data, paired with other drugs. There is no restriction on cell lines; the test set may include cell lines seen during training. LPO evaluates the model’s ability to predict interactions between drug pairs it has not encountered before. This is the most commonly used strategy in the literature.
- **Leave-Cell-Line-Out (LCO):** This strategy sets aside test data based on cell lines. Consequently, the train and test triplets do not share common cell lines. LCO tests the model’s ability to generalize predictions to new biological contexts represented by unseen cell lines. This is crucial for evaluating how well the model can adapt to different cellular contexts, which is important for applications like personalized medicine, where patient-specific cell responses are considered.
- **Leave-One-Drug-Out (LODO):** In stratification, at least one drug in each drug pair within the test set is completely absent in the training triplets. The other drug in the pair may still appear in the training set, interacting with different drugs. There is no restriction on cell lines; the test set may contain cell lines seen during training. The purpose of LODO is to evaluate the model’s ability to predict interactions when it has incomplete information—specifically when one drug is entirely new to the model. This scenario reflects real-world situations where a new drug is introduced.
- **Leave-Drug-Out (LDO):** If a drug is seen in the training data, this drug and all of its drug interactions are excluded from the test set. As a result, the model does not see any of the test drugs during training. Implementing this split requires first splitting the drugs and then adding their associated triplet; this reduces the number of training examples (Figure 1). There is no restriction on cell lines; the test set can include cell lines in the training data. LDO evaluates how well it can predict interactions involving drugs it has never seen in the training.

### Experimental Setup

In our experiments, we adopted the hyperparameters specified in the original papers for each model. For dataset splits, we utilized the original splits provided by the authors. Regarding DeepSynergy, we conducted LPO 5-fold nested cross-validation, consistent with the procedure described in the original paper, using the folds provided by the authors. For DeepDDS, we followed the LTO split procedure outlined in the original work, applying 5-fold cross-validation. In the MARSY experiments, the model employed the LPO split method along with 5-fold cross-validation. For JointSyn, the LTO split was generated using the authors’ provided code, resulting in 10 replicates per fold.

While the MatchMaker paper initially utilized the LPO split, we applied additional split strategies to study generalization and the behaviour of one-hot encoded features under different conditions. These included LPO, LCO, LODO, and LDO. In each case, 60% of the triplets are set for training, 20% for validation, and 20% for testing. For each split strategy, we generated 10 different train/test splits by varying the random split state. Results averaged across these splits are reported. Due to the nature of the LDO split, where the test set contains only unseen drugs, not all data could be utilized. Consequently, the DrugComb and NCI-ALMANAC datasets were reduced by approximately 68%

All models were trained using both their original drug and cell line features as well as one-hot-encoded features constructed based on their datasets. For DeepDDS and JointSyn, when training one-hot-encoded feature models, the graph components of the original architectures were removed, and one-hot-encoded features were directly provided as input to the models. The original architecture of DeepDDS and JointSyn, including the graph components, was fully maintained using its drug and cell line features.

### Synthetic Data Experiments

To assess models’ reliance on true features under different settings, we conducted synthetic data experiments. We constructed two datasets of drug–drug–cell line triplets: i) a *non-repeated* set of 24,500 unique (Drug_1_, Drug_2_, CellLine) combinations, where each drug and cell line is unique ii) a *repeated* set formed by fully crossing a panel of 50 drugs with 50 cell lines, yielding 61,250 triplets where drugs or cell lines can repeat in different combinations.

#### Synthetic setup 1: linear synergy score model

For each drug and cell line we created synthetic feature vectors by sampling from a standard normal distribution: each drug was assigned to **x** ∈ ℝ^100^ ∼ 𝒩 (0, **I**), and each cell line was assigned to**z** ∈ ℝ^100^ ∼ 𝒩 (0, **I**). For every entity, exactly 20 features were designated *informative*; the remaining 80 features were non-informative (weight 0). Informative features were assigned feature weights sampled uniformly from [0.5, 10.0].

#### Synergy scores were generated by the linear model

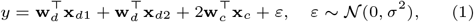

where **x**_*d*1_ and **x**_*d*2_ are the feature vectors of the two drugs, **x**_*c*_ is the context (cell line) feature vector, **w**_*d*_ and **w**_*c*_ are the corresponding weight vectors, and *ε* is Gaussian noise.

For prediction, we trained a linear regression model in PyTorch on concatenated inputs [**x**_*d*1_; **x**_*d*2_; **z**_*c*_] ∈ ℝ^300^ with MSE loss and L1 regularization at λ ∈ {10^−1^, 10^−2^, 10^−3^}. Training used early stopping on validation loss (max 1000 epochs). We trained models for the repeated dataset under the four split scenarios, each with 10 independent 60/20/20 train/val/test partitions. For the non-repeated dataset, we used 10 independent 60/20/20 splits without the need for different split types due to the absence of repetition.

#### Synthetic setup 2: nonlinear synergy score model

For a nonlinear synergy model, we generated synergy scores with a fixed three-layer MLP (256 → 128 → 1, ReLU) applied to masked, concatenated entity vectors. Specifically, each drug had **x** ∈ ℝ^150^ and each cell line **z** ∈ ℝ^200^ from 𝒩 (0, **I**); for every entity, 50 coordinates were designated informative and the rest set to zero. For each triplet, we concatenated 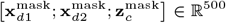, passed it through the fixed MLP (weight assigned uniformly at random), and scaled the output by 200 to obtain the synthetic synergy score. Scores were generated for both the non-repeating and repeating datasets described above.

We then trained a predictive MLP regressor on the *full*, unmasked inputs in ℝ^500^ (two drugs and one cell line), using architecture 512 → 128 → 1 with ReLU and dropout, MSE loss, early stopping, and L1 penalties λ ∈ {10^−1^, 10^−2^, 10^−3^}. The same split protocols were used as in the linear setup.

#### Assessment of Feature Importance and Feature Recovery

Feature importance for the linear model was quantified using the absolute value of the standardized coefficients. For the non-linear model, we applied Integrated Gradients (IG)(Sundararajan *et al*., 2017) to compute feature attributions using Captum library(Kokhlikyan *et al*., 2020). IG was computed with a zero baseline and 100 integration steps on 10% of the test set in each fold and averaged attributions across folds. We measured alignment by the Jaccard index between the top-k IG-attributed coordinates (*k* = 20 for the linear model, *k* = 50 for the nonlinear model) per drug or cell line and the known informative set.

### Computational Environment

We performed DeepSynergy, MatchMaker, MARSY, JointSyn, and DeepDDS experiments on systems equipped with Tesla V100-PCIE-32GB and Tesla V100S-PCIE-32GB GPUs. The Tesla V100-PCIE-32GB featured 31.74 GiB of memory, 80 cores, and a memory bandwidth of 836.37 GiB/s, while the Tesla V100S-PCIE-32GB offered the same memory and cores but with a bandwidth of 1.03 TiB/s. These systems included CPUs with frequencies ranging from 2.29 GHz to 3.59 GHz. TensorFlow with CUDA 10.1 and cuDNN 7 was used for training.

## Results

To assess the utility of the biochemical information, for each selected model, we trained it using its original biological and chemical features and compared it with a version where these features were replaced by one-hot encodings. By stripping away the biological and chemical information, we aim to assess to what extent the models rely on their intended features. We evaluated each model with its original source, dataset, and train/test splits. The details of these setups are provided in Section Methods, and the code reproducing the analysis is available at https://github.com/tastanlab/ohe.

### Comparison of the Models with the OHE Baseline Models

Figure 2 shows that the performance differences between the original drug and cell line features and OHE representations are minimal across all the models tested. For MatchMaker on the DrugComb dataset under the LPO split, the MSE difference between the original features and OHE representations was just 3.11%, indicating that the model performance remains largely unaffected if the drug and cell line molecular features are removed. DeepSynergy exhibited a minimal deviation (−0.91%) on the O’Neil dataset with OHE representations, demonstrating comparable performance to results obtained using the original drug and cell line features. Notably, MARSY, tested on the DrugComb dataset, showed a marginal improvement (−6.4%) when OHE features were used instead of the original feature. It suggests that the model’s predictive performance does not depend on the chemical and biological features. The performance gap was negligible for models like JointSyn and DeepDDS, which incorporate graph-based embedding techniques. JointSyn reported only a 0.59% deviation in MSE under the LTO split, while DeepDDS showed a mere 1.08% deviation in AUC. Thus, the results were persistent even for more complex graph-based architectures.

**Fig. 2.**
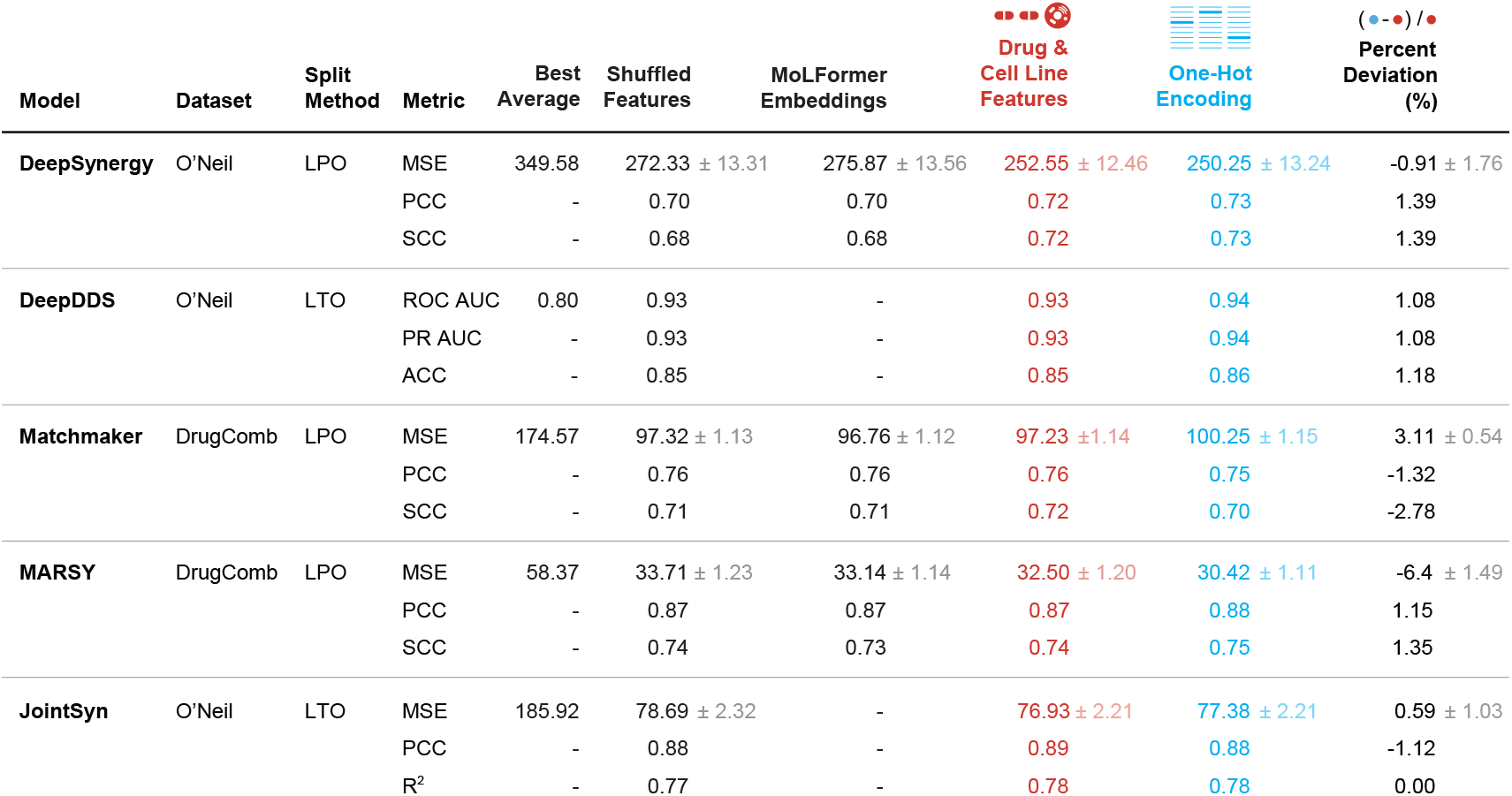
Comparison of Best Average Baselines, Shuffled Features, MoLFormer Embeddings, Drug & Cell Line Features, and OHE Representations Across Different Models and Datasets. The Best Average baseline reports the best result among four simple average-based predictors: overall average, drug pair average, cell line average, and cell line & at least one drug average. In the shuffled features setting, the feature distribution is preserved while the association with specific drugs or cell lines is broken. In the MoLFormer embeddings setting, drugs are represented by embeddings obtained from a pretrained MoLFormer model, while cell lines are represented with one-hot vectors. Drug and cell line features refer to the original representations used in the respective models. Performance metrics, percent deviation, and standard error (SE) are included in the table. The details of the SE calculations are provided in the Supplementary Section on Standard Error Calculations. For clarity, SE values less than 0.01 are omitted from the table but are available in Supplementary Table S5.

The performances of the models remain largely consistent regardless of whether original features or OHE representations were used. This consistent finding across a different set of synergy prediction models points out that the performance does not genuinely depend on the intended biological or chemical information provided to the model. Instead, the models take shortcuts by learning from the covariation patterns of the measurements. This could be an instance of shortcut learning (Geirhos *et al*., 2020).

We also compare these results to the average baseline predictor. For the average predictor baseline, the best performing average baseline is listed in Figure 2 while the complete results are in Supplementary Table S7. The results show that models using biological drug and cell line features perform significantly better than the strongest average-based baselines in most scenarios. In particular, under easier splitting strategies such as LPO, the performance gap is notable (with percent deviations ranging from 38% to 141%) (See Supplementary Table S7), indicating that the models are able to learn meaningful patterns from the training data. However, in more challenging split strategies (especially LDO and LCO), some models perform worse than even the best average baseline.

We also evaluated two additional controls (Figure 2): shuffled features and pretrained MoLFormer embeddings. The results were also consistently close or, in some cases, a little bit worse than the one-hot-encoded embeddings.

### Learning Residual of OHE Models

To assess whether the biological features contain information beyond what is captured by covariation patterns of the target values, we conducted an additional experiment, where we trained a model to predict the residuals of the OHE trained model using the biological features. DeepDDS was excluded from this analysis, as it is a classification model. Once the residual models were trained, we combined them with the corresponding OHE models to form an ensemble for the final prediction: *ŷ*_final_ = *ŷ*_OHE_ + *ŷ*_residual_

We evaluated whether the ensemble model could outperform either the original model trained on biological features or the OHE model. Across architectures, the ensemble’s performance was comparable to both with no measurable improvement (Supplementary Table S8). In most cases, the residual trained models failed to capture the OHE residuals, as indicated by near-zero correlations between predicted and true residuals —JointSyn being the sole exception. However, even for JointSyn, combining predictions with the OHE model did not enhance performance. Overall, these findings confirm that the biological features did not provide additional predictive value beyond what is captured from the covariation patterns.

### A Deeper Analysis with MatchMaker

To investigate generalization further, we experimented with MatchMaker in other evaluation setups that use different split strategies (explained in Methods). We omit the LTO strategy as this is the most straightforward scenario, and the models generally perform very well. We compared MatchMaker’s performance, which uses chemical descriptors and gene expression levels, with the model trained on the OHE representations for drugs and cell lines. We conducted these experiments both on DrugComb and NCI-ALMANAC datasets.

The model architectures correctly exploit underlying covariation patterns in the synergy measurements to make predictions for drug combinations within a tested drug panel and cell line, as indicated by the LPO split. However, the results degrade as the evaluation strategy sets a harder challenge. The models’ performance worsens on unseen cell lines, as shown in the LCO results. The results obtained on LODO, when one of the drugs is not seen, are even lower. The model fails to generalize to the case when both drugs in the test data are new, the LDO setup. These results show the models’ inability to transfer this knowledge to new, unseen drugs and cell lines.

The models trained with original features and OHE features show comparable performance on the LPO and LTO splits. The models relying on OHE struggle significantly when predicting synergy for drug pairs not present in the training set, the LDO scenario. In OHE, each drug is treated as an independent category without shared features that facilitate extrapolation to unseen combinations. Models utilizing feature-rich representations potentially encode drug similarities and relationships, enabling better generalization. Nevertheless, the results are still poor for both models in the LDO case.

### The Repeat Structure of the Data Leads to Shortcutting of the True Features

We hypothesize that models shortcut by exploiting the triplet structure of the data: when drugs or cell lines recur in different combinations, even if no exact drug–cell–dose triplet repeats, their feature vectors function as identifiers. In this view, prediction reduces to a tensor-completion problem over two drugs and a cell line, with missing synergy scores imputed from label covariation rather than biochemical signal. To study this under controlled conditions, we constructed two different datasets with different structures: a *repeating dataset*, where drugs and cell lines may recur (but exact triplets do not), and a *non-repeating dataset*, where no entity repeats. In practice, constructing a truly zero-shot triplet set, where the members of the triplets are never seen in the training, is not possible with the current datasets. Our synthetic experiments allowed us to construct a test set that is completely dissimilar to train drugs and cell lines. In the repeating dataset scenario, we experimented in all four split scenarios LPO, LCO, LODO, and LDO. The repeating dataset scenario inherently includes only unseen drugs and cell lines since no drug or cell line repeats in any of the triplets. We assessed whether models recover the designated ground-truth features. We then trained predictors with varying L1 regularization and evaluated models’ reliance on the true ground-truth features.

#### Linear synthetic synergy model results

On the repeating dataset, the linear predictor attains near-perfect accuracy under LPO but degrades when generalization requires unseen entities (LCO, LODO, LDO). For example, LPO yields MSE ≈ 0.01 and PCC/SCC = 1.00 across λ values, whereas LCO and LDO show much larger errors despite reasonably high correlations (Supplementary Table S9). In contrast, on the non-repeating dataset, the same model achieves near-perfect performance for all λ (Supplementary Table S10).

What is more interesting is the difference in the reliance on the true features. Under repeated splits (LPO), weak regularization (λ = 10^−3^) produces many non-zero coefficients on irrelevant features, and for LCO, LODO, and LDO even strong regularization still yields many non-zero coefficients (Supplementary Figures S3–S6). Strikingly, models trained on the non-repeating dataset yield very clean recovery of the informative features (Supplementary Figure S7). Figure 3 exemplifies this difference in the reliance of features for the LDO and the LCO setups under different regularization penalties. The strong recovery in the non-repeating setup persists even when the training set is subsampled to 10% of the original dataset size (Supplementary Figure S8). Thus, in the non-repeated setting, models can learn to depend on the set of informative features, which help them generalize to unseen drugs and cell lines. These results indicate that repetition encourages shortcut solutions that learn from covariation, whereas less repetition drives the model to base its learning on the true features.

**Fig. 3.**
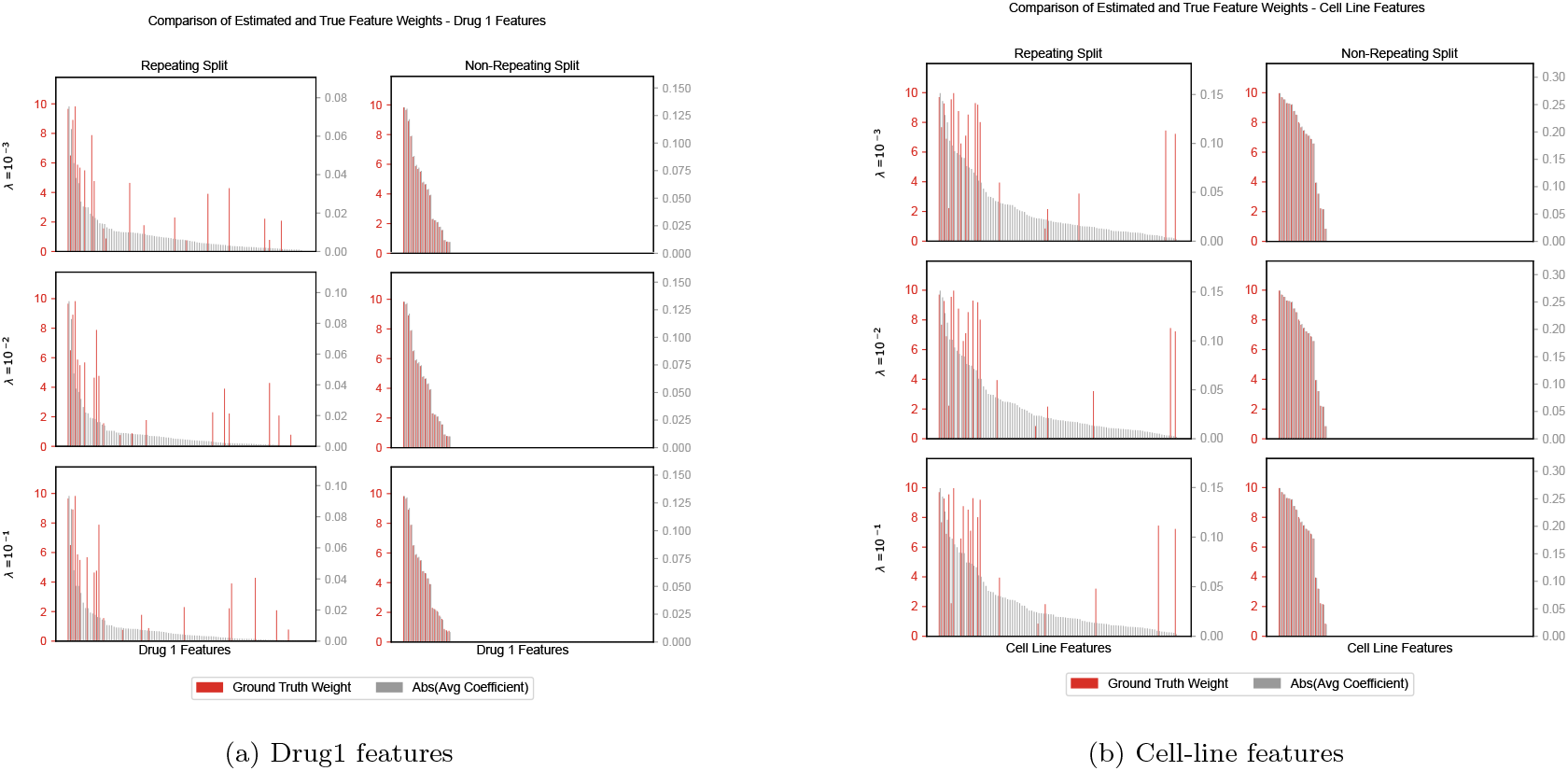
Red bars show the ground-truth weights and gray bars show the average linear-model coefficients. (a) For the drug features, the non-repeating split makes the model align with the informative features, whereas the repeating split(LDO) leads to large weights on non-informative ones. (b) For the cell-line features, the non-repeating split again highlights the informative signals, while the repeating split(LCO) causes spurious weights on irrelevant features.

#### Nonlinear synthetic synergy model results

In the nonlinear synergy score setup, we observe the same behavior. On the repeating dataset, predictive performance is moderate to high but varies by split and λ (e.g., PCC ≈ 0.45−0.85 (Supplementary Table S11). On the non-repeating dataset, performance is stable (PCC/SCC ≈ 0.67) across λ (Supplementary Table S12). Under the repeating dataset learning, the feature attributions show limited alignment with ground-truth features in LPO (Supplementary Figure S9) and also in other split settings (shown in Supplementary Figures S10-S12). We find that the feature attribution distributions between the relevant and irrelevant features differ only in the non-repeating setting (Supplementary Figure S13). Thus, the non-repeating setting drives the model to strongly rely on the true features. The Jaccard indices between the set of high attribution feature set and the ground-truth feature set results corroborate this: the overlap of the ground truth feature sets and the important feature sets are low for the models trained on the repeating dataset and substantially higher in the non-repeated dataset. The Jaccard indices are summarized in Table 5.

**Table 4.** Performance Comparison of MatchMaker Model Using Drug & Cell Line Features vs OHE Representations on NCI-ALMANAC Dataset Across Different Split Methods. Chemical descriptors were originally used to represent drug features, while cell line gene expression levels represented cell line features in the MatchMaker model. These representations serve as the baseline for comparison with one-hot-encoded drug and cell line representations.

| Split Method | Feature Type | MSE | PCC | SCC |
| --- | --- | --- | --- | --- |
| LPO | Drug & Cell Feature | 2912.40 ± 68.03 | 0.57 ± 0.012 | 0.54 ± 0.005 |
| | One-Hot-Encoding | 3041.02 ± 70.18 | 0.54 ± 0.010 | 0.52 ± 0.002 |
| LCO | Drug & Cell Feature | 2956.30 ± 43.27 | 0.57 ± 0.007 | 0.46 ± 0.005 |
| | One-Hot-Encoding | 2930.47 ± 45.97 | 0.57 ± 0.006 | 0.47 ± 0.005 |
| LODO | Drug & Cell Feature | 3903.41 ± 83.75 | 0.26 ± 0.014 | 0.27 ± 0.012 |
| | One-Hot-Encoding | 3943.30 ± 84.28 | 0.24 ± 0.010 | 0.25 ± 0.004 |
| LDO | Drug & Cell Feature | 4587.08 ± 152.46 | 0.16 ± 0.021 | 0.17 ± 0.018 |
| | One-Hot-Encoding | 4670.97 ± 154.58 | 0.09 ± 0.010 | 0.09 ± 0.007 |

**Table 5.**
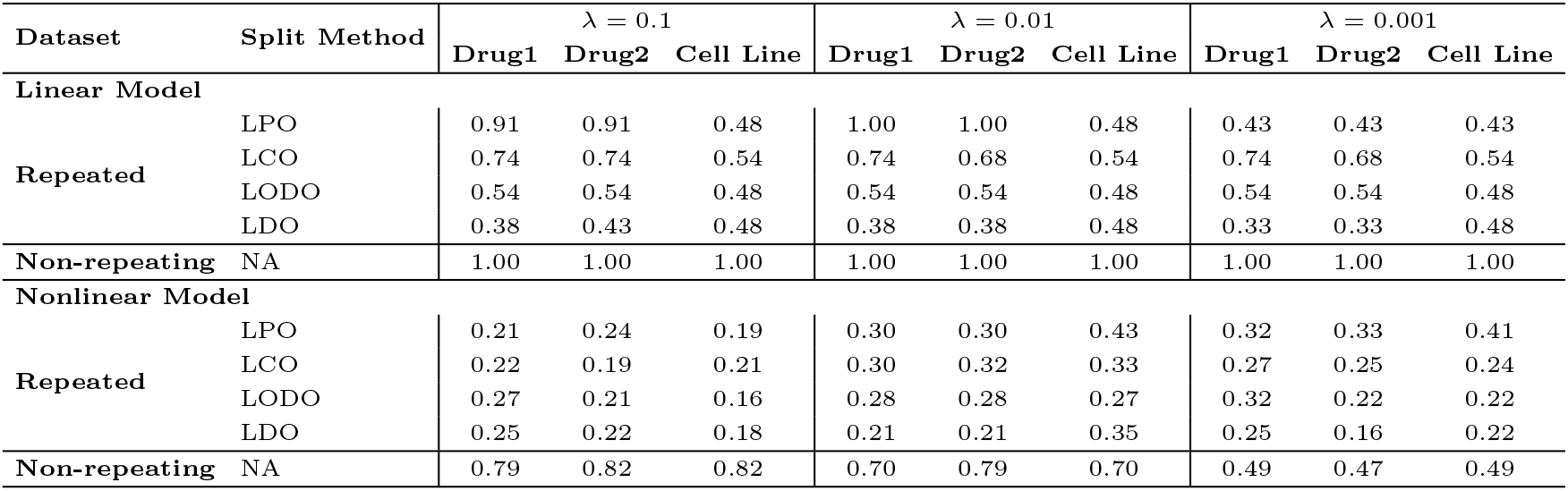
Feature recovery success of the model for the linear and the nonlinear synthetic synergy model setups. Jaccard overlap between model-identified features and ground-truth informative features on the *repeated* and *non-repeated* datasets. Columns report Jaccard index for Drug 1, Drug 2, and Cell Line across L1 penalties (λ). For the linear block, top-20 features are taken from standardized absolute coefficients (averaged over 10 random splits). For the nonlinear block, top-50 features are taken from Integrated Gradients (IG). *NA* indicates “not applicable”: in the non-repeated dataset, no entity recurs, therefore LPO/LCO/LODO/LDO are undefined.

### On the relationship of training counts and error

To check whether training counts can explain model errors, we carried out analyses on the DrugComb and MatchMaker; for each split type and fold, we fit a linear model:

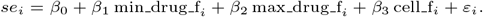

where the predictors are the minimum and the maximum training frequencies of the two drugs (min drug f, min drug f and the frequency of the cell line(cell f). Across all split–fold combinations, the *R*^2^ values range between 0.000 and 0.010, meaning that these count variables explain at most about 1% of the variance in squared error. This suggests that training counts, by themselves, are very weak predictors of model error.

We also examined the error as a function of pair-level counts. For LPO, LCO, and LODO, we additionally trained shallow regression trees (maximum depth 2) using binned frequency features to predict the error of a test triplet based on the training frequencies of its constituent entities. The aim here is to check if these counts explain the error observed across triplets. For LPO, we used the minimum train+validation frequency across the two drug–cell pairs (drug1–cell and drug2–cell). For LCO, we used the drug-pair count, and for LODO we used the maximum drug–cell frequency. All count features were binned into the intervals {0, 1, 2–99, 100–999, 1000–2999, ≥ 3000}. These trees are shown in Supplementary Figures S14-16.

Overall, the trees show that very rare pairs do tend to have higher errors, but this is expected of any prediction task. Thus, pair-level frequencies capture only part of the effect and cannot fully explain the model’s error patterns. These analyses suggest that the model performances cannot be explained by per-drug or per–cell line frequency counts.

## Discussion

Effective predictions can streamline the identification of effective drug combinations in different biological contexts, reducing the time and cost associated with experimental validation. Although current synergy prediction models report high predictive performance when tested on new combinations of drugs or cell lines observed in training, they exhibit significant limitations in generalizing to unseen drugs. In this work, by replacing the biological and chemical information on drugs and cell lines with simple identifiers, we show that the models learn from the covariation patterns of the synergy measurements rather than from the domain-relevant features.

Our objective was not to compare different architectures and find the best one or to identify the most effective feature encodings—readers interested in such benchmarks are referred to (Abbasi and Rousu, 2024; Tasnina *et al*., 2025). Rather, we sought to determine whether each model truly leverages its intended input features. To this end, we held all other factors constant—using identical data splits, hyperparameter settings, and training protocols—and replaced the original drug and cell-line descriptors with simple one-hot encodings. By evaluating model performance under these controlled conditions, we could directly assess the extent to which models depend on biologically informed feature representations. This study once again highlights that diagnosing model behaviors with different strategies is imperative for understanding the capabilities and advancing these models. These results also caution us to have multiple baseline models to interrogate the predictions.

MARSY, along with synergy scores, predicts single-drug responses. In our experiment, we also observed a similar result for single drug response prediction in MARSY; models trained with OHE, despite lacking any transcriptomic information, perform on par with the models trained with the biological features (Supplementary Table S6). This finding suggests that this issue may extend beyond the drug synergy prediction domain and might be prevalent for single-drug response predictions as well. Indeed, a recent study reports a significant performance drop for drug response prediction when models are tested on unseen datasets (Partin *et al*., 2026). Other studies also report instances where models fail to generalize to novel chemistry, such as in drug-target interaction prediction (Luo *et al*., 2024; Vefghi et al., 2025; Wang et al., 2023a; Chatterjee *et al*., 2023; Ong et al., 2023), bioactivity prediction (Theisen *et al*., 2024), and de novo molecule design (Nori and Jin, 2024; Shukueian Tabrizi *et al*., 2025; Albrijawi and Alhajj, 2024).

In this drug synergy prediction task, our results show that the true features are bypassed. Shortcut learning instances have been reported in the literature for deep vision and natural language processing tasks (Geirhos *et al*., 2020; Hermann et al., 2023). For example, a deep neural network might seem to recognize cows well, but it struggles with images where cows are not set against a typical grassy background, exposing that it had inadvertently used the presence of grass as a shortcut to predict cows (Beery *et al*., 2018). Other well-known examples include a model learning to use the presence of rulers in lesion images as a shortcut for predicting malignancy (Narla *et al*., 2018) or learning based on the presence of hospital identifiers in X-ray scans (Zech *et al*., 2018; DeGrave *et al*., 2021). These shortcuts are caused by confounding correlations between the inputs and outputs. In this case, the shortcut is different. There is no confounding variable to shortcut; the model bypasses the true features using the feature vectors and relies only on the covariation in the target synergy values. Using controlled synthetic datasets, we show that repeating drugs and cell lines push models to exploit label covariation tied to drug or cell line identity rather than the intended features. When repetition is removed, predictors can perfectly recover the designated informative features with high fidelity. These results implicate the triplet formulation itself as a driver of shortcut learning. Notably, many knowledge-graph tasks share an analogous (head, relation, tail) triplet structure with heavy entity reuse. Generalization issues in knowledge graph link prediction tasks have been reported (Brière *et al*., 2025). It would be an interesting research direction to investigate whether the generalization issues are due to the repeating entity structure.

Despite the challenges in generalizing to novel drug pairs, models that capture covariation patterns remain valuable for discovering new synergistic combinations within an existing drug panel. By leveraging the learned interactions and correlations, these models can efficiently identify promising combinations that warrant further experimental testing, thereby accelerating the drug discovery process within the scope of the tested drugs. However, broader generalization remains an area for improvement.

## Author contributions statement

B.C. collected data, conducted the experiments, and wrote the first draft of the manuscript; H.İ.K. contributed to software development and reviewed the manuscript; A.E.C. and M.R. analyzed the results and reviewed the manuscript; O.T. conceived the experiments, analyzed results, and wrote and reviewed the manuscript.

