## Supplementary File for "One-Hot News: Drug Synergy Models Shortcut Molecular Features"

### Supplementary Materials for "One-Hot News: Drug Synergy Predictors Shortcut Molecular Features"

#### 1 Chemical Features of MatchMaker

The chemical features of drugs are derived using the PyBioMed library. These descriptors include various chemical and physical properties of the molecules. 12 distinct descriptor categories, comprising 367 individual features, are computed using the available functionalities in the library. Table S1 provides a detailed breakdown of these descriptor categories and their corresponding feature counts.

Table S1: Drug chemical structure features collected from PyBioMed.

| **Descriptor Category** | **Count** |
|---|---|
| Kappa Descriptors | 7 |
| Charge Descriptors | 25 |
| Connectivity Descriptors | 44 |
| Constitution Descriptors | 28 |
| Geary Descriptors | 32 |
| MOE Descriptors | 59 |
| Moran Descriptors | 32 |
| Moreau-Broto Descriptors | 32 |
| Topology Descriptors | 19 |
| Molecular Properties | 4 |
| Basak Descriptors | 21 |
| Burden Descriptors | 64 |

### 2 MatchMaker Dataset Statistics

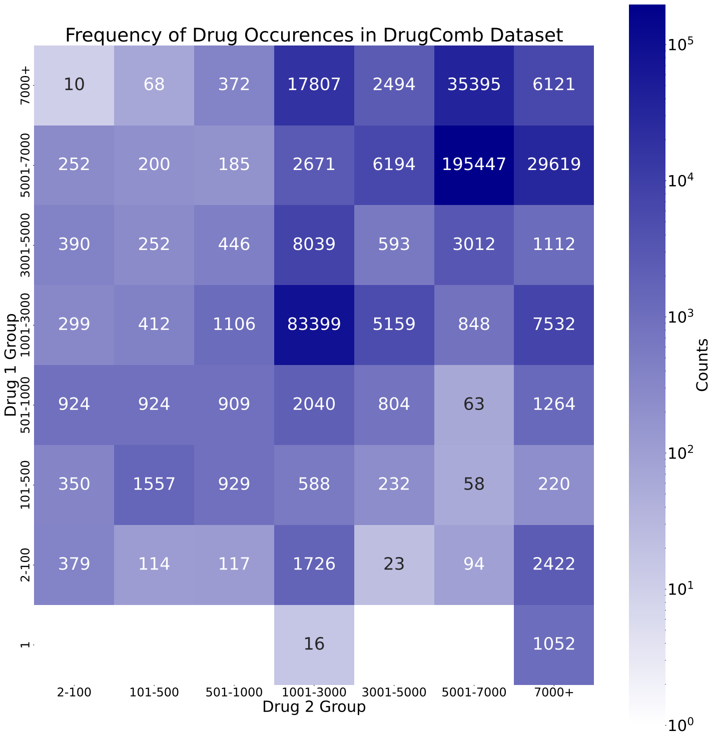


Figure S1: Heatmap representing the frequency of drug occurrences and their pairwise groupings in the DrugComb dataset. Drug frequencies are categorized into distinct groups based on their occurrence counts in triplets.

Table S2: Total samples for 10 splits using LDO split strategy on DrugComb dataset.

| **Repeat** | **Total Samples** |
|---|---|
| Repeat 1 | 150,055 |
| Repeat 2 | 150,864 |
| Repeat 3 | 149,532 |
| Repeat 4 | 151,050 |
| Repeat 5 | 152,334 |
| Repeat 6 | 152,903 |
| Repeat 7 | 150,468 |
| Repeat 8 | 149,513 |
| Repeat 9 | 151,725 |
| Repeat 10 | 154,461 |

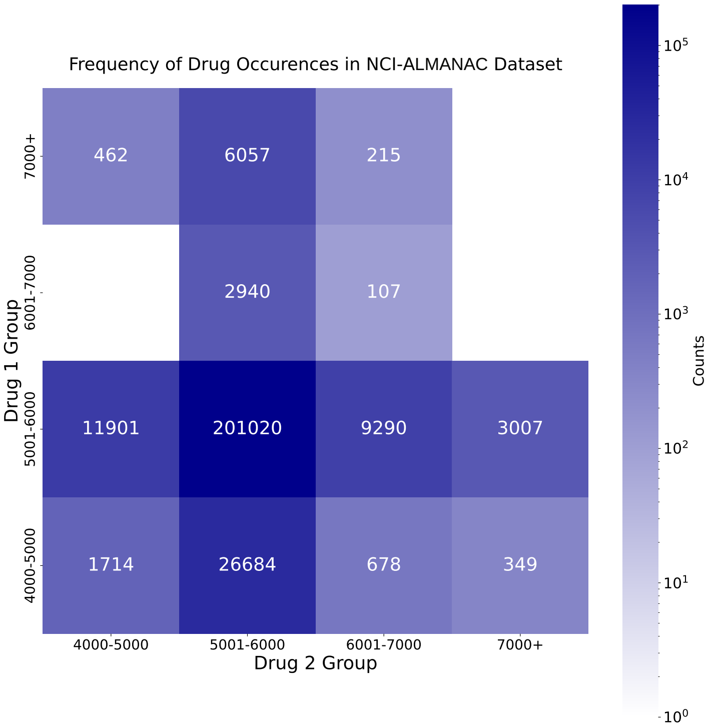


Figure S2: Heatmap representing the frequency of drug occurrences and their pairwise groupings in the NCI-ALMANAC dataset. Drug frequencies are categorized into distinct groups based on their occurrence counts in triplets.

Table S3: Total samples for 10 splits using LDO split strategy on NCI-ALMANAC dataset.

| **Repeat** | **Total Samples** |
|---|---|
| Repeat 1 | 93,139 |
| Repeat 2 | 92,831 |
| Repeat 3 | 94,223 |
| Repeat 4 | 92,215 |
| Repeat 5 | 92,973 |
| Repeat 6 | 92,529 |
| Repeat 7 | 94,169 |
| Repeat 8 | 94,842 |
| Repeat 9 | 93,271 |
| Repeat 10 | 92,287 |

### 3 Performance Comparisons of Original Features vs. OHE Across Models

Table S4 provides the repositories where the model source codes are obtained.

Table S4: Drug Synergy Prediction Models and Their GitHub Repositories

| **Model** | **GitHub Source** |
|---|---|
| MatchMaker | `https://github.com/tastanlab/matchmaker/tree/master` |
| DeepSynergy | `https://github.com/KristinaPreuer/DeepSynergy/tree/master` |
| MARSY | `https://github.com/Emad-COMBINE-lab/MARSY` |
| JointSyn | `https://github.com/LiHongCSBLab/JointSyn` |
| DeepDDS | `https://github.com/Sinwang404/DeepDDs/tree/master` |

Table S5: Comparison of Drug & Cell Line Features, One-Hot Encoding(OHE), Shuffled Features, and MoLFormer Embeddings Across Different Models and Datasets. Detailed performance metrics and standard error (SE) values, including those omitted from the main table ($SE < 0.01$), are presented. Drug and cell line features represent the original input representations used in the models. Percent deviation and SE calculation methods are described in the Standard Error Calculations Section. Metrics reported: MSE (mean squared error), PCC (Pearson correlation coefficient), and SCC (Spearman correlation coefficient).

| **Model** | **Dataset** | **Split Method** | **Metric** | **Drug & Cell Line Features** | **One-Hot Encoding** | **Shuffled Features** | **MoLFormer Embeddings** | **Percent Deviation (%)** |
|---|---|---|---|---|---|---|---|---|
| DeepSynergy | O'Neil | LPO | MSE | 252.55 ± 12.46 | 250.25 ± 13.24 | 272.33 ± 13.31 | 275.87 ± 13.56 | -0.91 ± 1.76 |
| | | | PCC | 0.72 ± 0.005 | 0.73 ± 0.008 | 0.70 ± 0.008 | 0.70 ± 0.007 | 1.39 ± 0.006 |
| | | | SCC | 0.72 ± 0.004 | 0.73 ± 0.005 | 0.68 ± 0.003 | 0.68 ± 0.004 | 1.39 ± 0.003 |
| DeepDDS | O'Neil | LTO | ROC AUC | 0.93 ± 0.003 | 0.94 ± 0.001 | 0.93 | - | 1.08 ± 0.003 |
| | | | PR AUC | 0.93 ± 0.004 | 0.94 ± 0.002 | 0.93 | - | 1.08 ± 0.003 |
| | | | ACC | 0.85 ± 0.006 | 0.86 ± 0.004 | 0.85 | - | 1.18 ± 0.005 |
| MatchMaker | DrugComb | LPO | MSE | 97.23 ± 1.14 | 100.25 ± 1.15 | 97.32 ± 1.13 | 96.76 ± 1.12 | 3.11 ± 0.54 |
| | | | PCC | 0.76 ± 0.002 | 0.75 ± 0.002 | 0.76 ± 0.001 | 0.76 ± 0.001 | -1.32 ± 0.002 |
| | | | SCC | 0.72 ± 0.001 | 0.71 ± 0.002 | 0.71 ± 0.002 | 0.71 ± 0.001 | -2.78 ± 0.003 |
| MARSY | DrugComb | LPO | MSE | 32.50 ± 1.15 | 30.42 ± 1.11 | 33.71 ± 1.23 | 33.14 ± 1.14 | -6.40 ± 1.49 |
| | | | PCC | 0.87 ± 0.004 | 0.88 ± 0.003 | 0.87 ± 0.003 | 0.87 ± 0.004 | 1.15 ± 0.002 |
| | | | SCC | 0.74 ± 0.005 | 0.75 ± 0.005 | 0.74 ± 0.005 | 0.73 ± 0.005 | 1.35 ± 0.003 |
| JointSyn | O'Neil | LTO | MSE | 76.93 ± 2.21 | 77.38 ± 2.21 | 78.69 ± 2.32 | - | 0.59 ± 1.03 |
| | | | PCC | 0.89 ± 0.001 | 0.88 ± 0.001 | 0.88 ± 0.001 | - | -1.12 ± 0.001 |
| | | | $R^2$ | 0.78 ± 0.001 | 0.78 ± 0.001 | 0.77 ± 0.001 | - | 0.00 |

Table S6: MARSY single response prediction results from Drug Cell Line feature and OHE experiments. RS1 and RS2 represent relative inhibition responses for drugs in a pair.

| **Metric** | **Feature Type** | **MSE** | **PCC** | **SCC** |
|---|---|---|---|---|
| RS1 | Drug & Cell Line Features | 36.72 ± 2.02 | 0.94 ± 0.002 | 0.91 ± 0.001 |
| | One Hot Encoded | 34.45 ± 2.16 | 0.94 ± 0.001 | 0.92 ± 0.001 |
| RS2 | Drug & Cell Line Features | 37.25 ± 2.09 | 0.94 ± 0.002 | 0.91 ± 0.001 |
| | One Hot Encoded | 34.26 ± 2.21 | 0.94 ± 0.002 | 0.92 ± 0.002 |

### 4 Standard Error Calculations

#### 4.1 Standard Error of MSE

The standard error of the MSE ($SE_{\text{mse}}$) is calculated based on the standard deviation ($\sigma$) of the squared errors and the sample size $N$, using the formula:

$$SquarredError = (\text{truth} - \text{pred})^2 \quad (1)$$

$$SE_{\text{mse}} = \frac{\sigma_{\text{squared\_error}}}{\sqrt{N}} \quad (2)$$

#### 4.2 Standard Error of Other Metrics

The standard errors of the PCC ($SE_{\text{pcc}}$), SCC ($SE_{\text{scc}}$), $R^2$ ($SE_{R^2}$), ROC AUC ($SE_{\text{roc\_auc}}$), PR AUC ($SE_{\text{pr\_auc}}$), and ACC ($SE_{\text{acc}}$) were calculated based on the standard deviation ($\sigma_{\text{metric}}$) of the metrics across folds and the number of folds $N$. The formula used for these calculations is as follows:

$$SE_{\text{metric}} = \frac{\sigma_{\text{metric}}}{\sqrt{N}} \quad (3)$$

where ($\sigma_{\text{metric}}$) represents the standard deviation of the metric values obtained from each fold, and $N$ is the total number of folds.

### 5 Percent Deviation and Their Standard Errors Calculations

The percent deviation is calculated as:

$$\texttt{Percent Deviation} = \frac{y_1 - y_2}{y_2} \quad (4)$$

where $y_1$ is the result for OHE and $y_2$ is the reference(drug & cell line features).

#### 5.1 Standard Error of Percent Deviation for MSE

The standard error of percent deviation for MSE is calculated as follows:

1. Calculate the difference between the squared errors of the predictions from the two representations:

$$\Delta = (\text{pred}_1 - \text{truth})^2 - (\text{pred}_2 - \text{truth})^2 \quad (5)$$

2. Calculate the standard deviation of the differences ($\sigma_\Delta$), then divide it by the square root of the number of folds ($N$):

$$SE_\Delta = \frac{\sigma_\Delta}{\sqrt{N}} \quad (6)$$

3. Scale by the reference MSE to express as a percentage. Divide $SE_\Delta$ by the reference MSE value ($\text{MSE}_2$) and multiply by 100:

$$SE_{\text{percent deviation}} = \frac{SE_\Delta}{\text{MSE}_2} \times 100 \quad (7)$$

#### 5.2 Standard Error of Percent Deviation for Other Metrics

To compute the standard error of percent deviation for a metric:

1. Compute the difference between the metric values obtained using one-hot encoding ($\text{metric\_value}_1$) and the reference metric values from drug and cell line features ($\text{metric\_value}_2$):

$$\Delta = \text{metric\_value}_1 - \text{metric\_value}_2 \quad (8)$$

2. Calculate the standard deviation of the differences ($\sigma_\Delta$), then divide it by the square root of the number of folds ($N$):

$$SE_\Delta = \frac{\sigma_\Delta}{\sqrt{N}} \tag{9}$$

3. Scale by the mean reference metric value. Divide $SE_\Delta$ by the mean reference metric value (metric_value$_2$) and multiply by 100 to express it as a percentage:

$$SE_{\text{percent deviation}} = \frac{SE_\Delta}{\text{metric\_value}_2} \times 100 \tag{10}$$

### 6 Average Predictor Baselines

Table S7: Average predictive performance of baseline compared to the models trained with original features.

| Model | Dataset | Split Method | Metric | Overall Average | Drug Pair Average | Cell Line Average | Cell Line & ≥1 Drug Average | Drug&Cell Line Features | Percent Dev. |
|---|---|---|---|---|---|---|---|---|---|
| DeepSynergy | O'Neil | LPO | MSE | 525.69 | 525.69 | 515.47 | 349.58 | 252.55 | 38.42 |
| DeepDDS | O'Neil | LTO | AUC ROC | 0.50 | 0.79 | 0.56 | 0.80 | 0.94 | -14.89 |
| Matchmaker | DrugComb | LPO | MSE | 227.54 | 227.54 | 219.43 | 174.57 | 97.23 | 79.54 |
| Matchmaker | DrugComb | LCO | MSE | 216.66 | 147.08 | 216.66 | 216.66 | 161.68 | -9.03 |
| Matchmaker | DrugComb | LODO | MSE | 225.93 | 225.93 | 218.52 | 191.49 | 199.22 | -3.88 |
| Matchmaker | DrugComb | LDO | MSE | 224.54 | 224.54 | 219.92 | 224.54 | 237.42 | -7.37 |
| MARSY | DrugComb | LPO | MSE | 133.50 | 133.50 | 123.68 | 58.37 | 32.50 | 79.60 |
| JointSyn | O'Neil | LTO | MSE | 347.62 | 185.92 | 334.98 | 204.23 | 76.93 | 141.67 |

### 7 Residual Model Performance

The residual performance of the model, the ensemble model is compared to the models trained with OHE and the Drug&Cell Line features.

Table S8: Performance of models trained on original biological features to predict the residuals of OHE models. The OHE models are trained to predict synergy scores, while the residual models are trained to predict the OHE models' residuals. Drug and cell line features correspond to those used in each model's original publication. The ensemble model combines predictions from the OHE and residual models.

| **Model** | **Metric** | Residual | OHE Encoding | Ensemble | Drug&Cell Line Features |
|---|---|---|---|---|---|
| DeepSynergy | MSE | 272.27 | 250.25 | 273.13 | 252.55 |
| | PCC | 0.00 | 0.73 | 0.70 | 0.72 |
| | SCC | -0.04 | 0.73 | 0.69 | 0.72 |
| MatchMaker | MSE | 99.61 | 100.25 | 99.61 | 97.23 |
| | PCC | 0.08 | 0.74 | 0.75 | 0.76 |
| | SCC | 0.07 | 0.70 | 0.70 | 0.72 |
| MARSY | MSE | 37.18 | 30.41 | 37.18 | 32.50 |
| | PCC | -0.05 | 0.88 | 0.85 | 0.87 |
| | SCC | -0.06 | 0.75 | 0.71 | 0.74 |
| JointSyn | MSE | 54.38 | 77.38 | 73.90 | 76.93 |
| | PCC | 0.40 | 0.88 | 0.89 | 0.89 |
| | SCC | 0.37 | 0.89 | 0.90 | 0.89 |

### 8 Results on Synthetic Synergy Score Experiments

#### 8.1 Predictive Performance of the Models

Below, the predictive performance of the models trained in the synthetic experiments setups are provided in Tables S9, S10, S11 and S12.

Table S9: The linear synthetic synergy model on the repeated dataset setup. Predictive under different L1 penalties ($\lambda$) and evaluation splits.

| **Split Type** | $\lambda = 0.1$ | | | $\lambda = 0.01$ | | | $\lambda = 0.001$ | | |
|---|---|---|---|---|---|---|---|---|---|
| | **MSE** | **PCC** | **SCC** | **MSE** | **PCC** | **SCC** | **MSE** | **PCC** | **SCC** |
| LPO | 0.31 | 1.00 | 1.00 | 0.01 | 1.00 | 1.00 | 0.01 | 1.00 | 1.00 |
| LCO | 642.71 | 0.79 | 0.78 | 639.31 | 0.79 | 0.95 | 643.66 | 0.79 | 0.78 |
| LODO | 43.50 | 0.99 | 0.99 | 43.04 | 0.99 | 0.99 | 51.89 | 0.99 | 0.98 |
| LDO | 203.61 | 0.94 | 0.93 | 202.77 | 0.94 | 0.93 | 209.94 | 0.94 | 0.93 |

Table S10: The linear synthetic synergy model on the non-repeated dataset setup. Predictive under different L1 penalties ($\lambda$) and evaluation splits.

| **Split Type** | $\lambda = 0.1$ | | | $\lambda = 0.01$ | | | $\lambda = 0.001$ | | |
|---|---|---|---|---|---|---|---|---|---|
| | **MSE** | **PCC** | **SCC** | **MSE** | **PCC** | **SCC** | **MSE** | **PCC** | **SCC** |
| Non-repeated | 0.15 | 1.00 | 1.00 | 0.01 | 1.00 | 1.00 | 0.01 | 1.00 | 1.00 |

#### 8.2 Linear model feature recovery figures

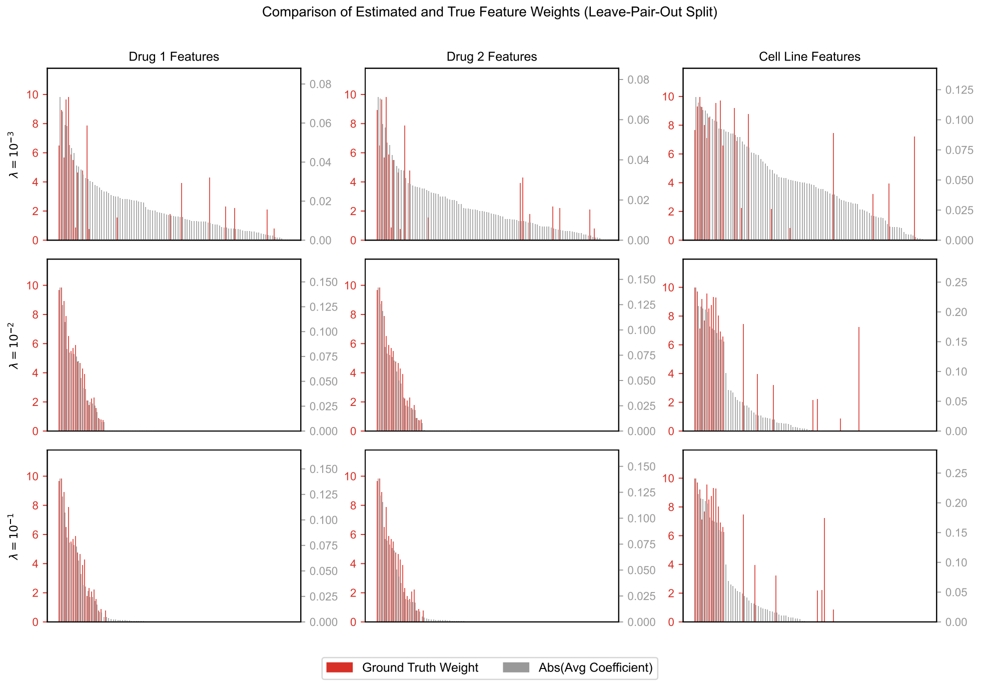


Figure S3: Absolute average standardized coefficients (gray bars) and ground-truth feature weights (red bars) for Drug 1, Drug 2, and cell-line features under the LPO split, with L1 regularization strengths $\lambda = 10^{-3}$, $\lambda = 10^{-2}$, and $\lambda = 10^{-1}$.

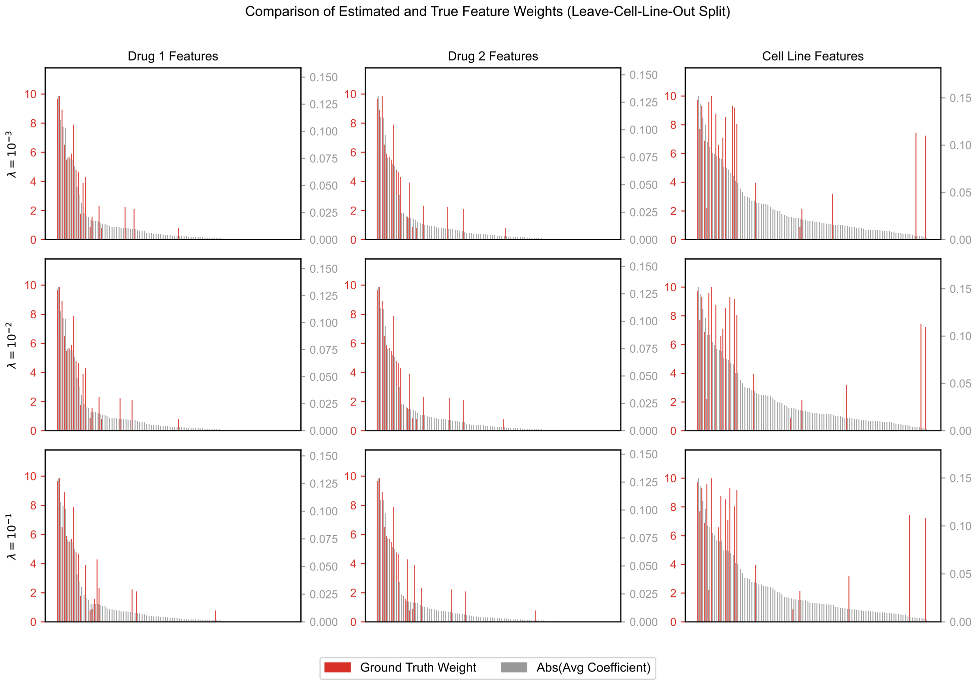


Figure S4: Absolute average standardized coefficients (gray bars) and ground-truth feature weights (red bars) for Drug 1, Drug 2, and cell-line features under the LCO split, with L1 regularization strengths $\lambda = 10^{-3}$, $\lambda = 10^{-2}$, and $\lambda = 10^{-1}$.

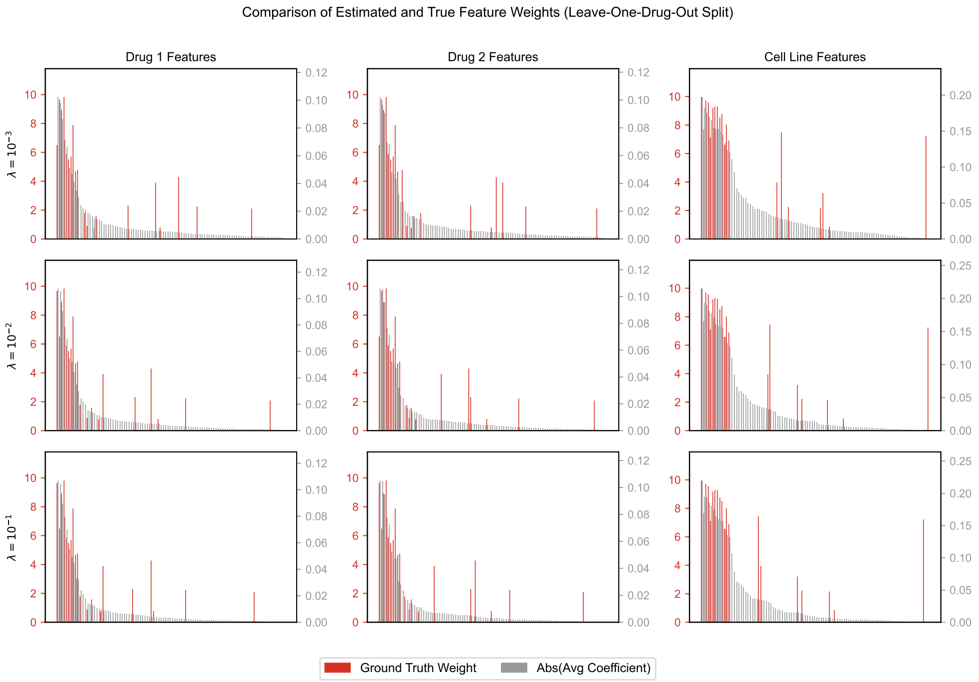


Figure S5: Absolute average standardized coefficients (gray bars) and ground-truth feature weights (red bars) for Drug 1, Drug 2, and cell-line features under the LODO split, with L1 regularization strengths $\lambda = 10^{-3}$, $\lambda = 10^{-2}$, and $\lambda = 10^{-1}$.

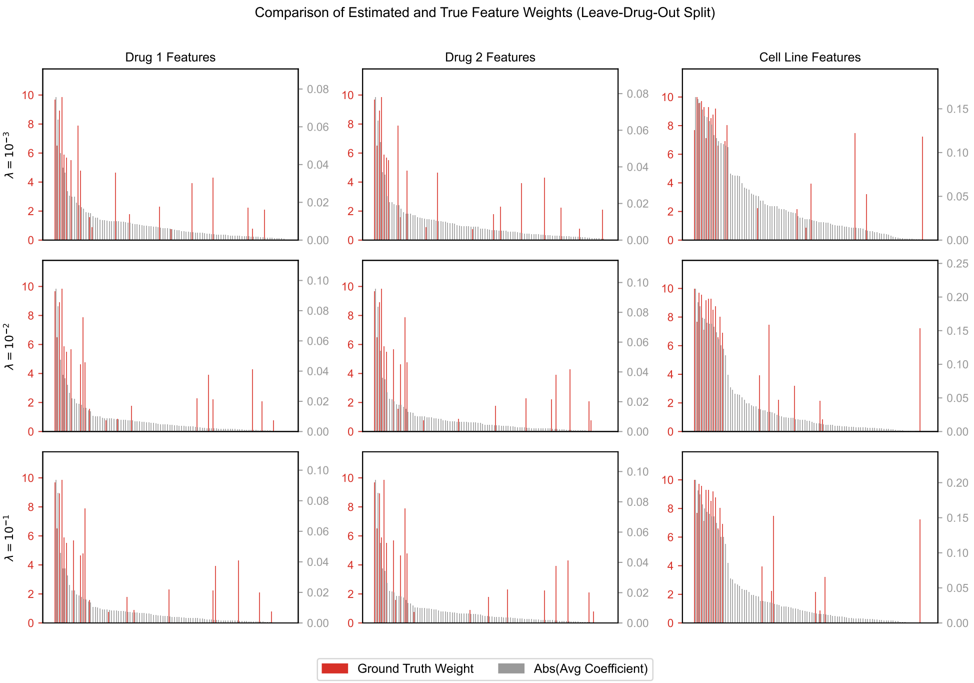


Figure S6: Absolute average standardized coefficients (gray bars) and ground-truth feature weights (red bars) for Drug 1, Drug 2, and cell-line features under the LDO split, with L1 regularization strengths $\lambda = 10^{-3}$, $\lambda = 10^{-2}$, and $\lambda = 10^{-1}$.

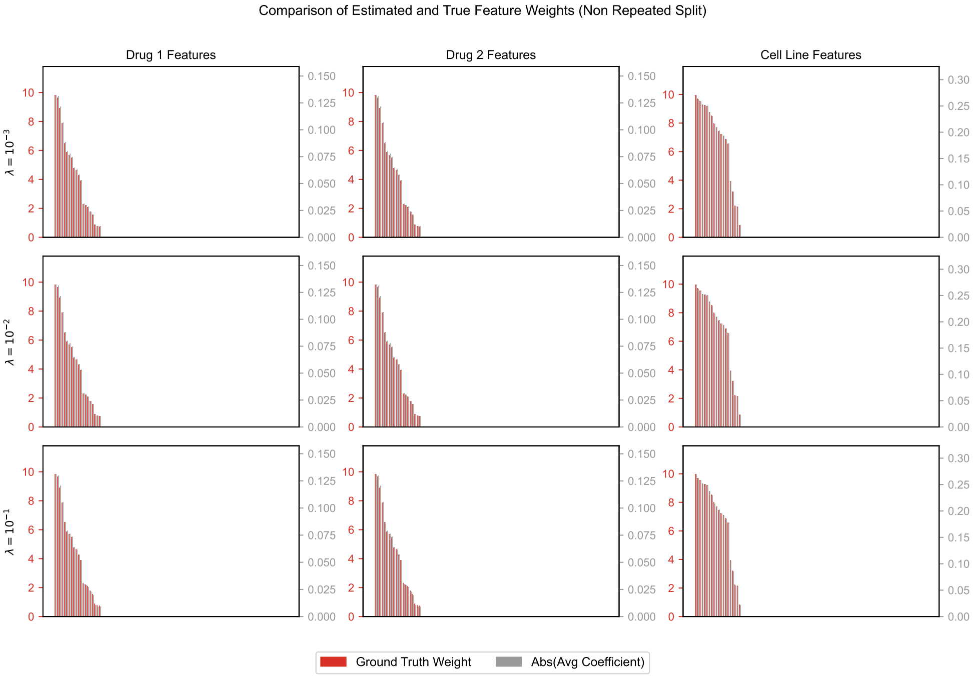


Figure S7: Absolute average standardized coefficients (gray bars) and ground-truth feature weights (red bars) for Drug 1, Drug 2, and cell-line features for the model trained and tested on non-repeating dataset $\lambda = 10^{-3}$, $\lambda = 10^{-2}$, and $\lambda = 10^{-1}$.

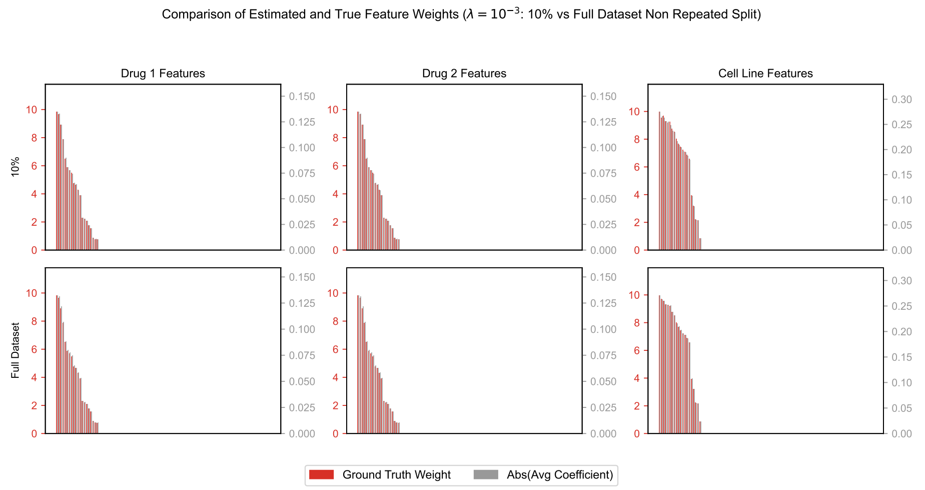


Figure S8: Comparison of learned feature importance (gray bars) and ground-truth weights (red bars) for Drug 1, Drug 2, and Cell Line features under two non-repeated settings: using the full dataset and using only 10% of the training set (both with $\lambda = 10^{-3}$).

#### 8.3 Nonlinear model feature recovery figures

Table S11: The non-linear synthetic synergy model on the repeated dataset setup. Predictive under different L1 penalties ($\lambda$) and evaluation splits.

| **Split Type** | $\lambda = 0.1$ | | | $\lambda = 0.01$ | | | $\lambda = 0.001$ | | |
|---|---|---|---|---|---|---|---|---|---|
| | **MSE** | **PCC** | **SCC** | **MSE** | **PCC** | **SCC** | **MSE** | **PCC** | **SCC** |
| LPO | 22.80 | 0.73 | 0.73 | 15.39 | 0.82 | 0.81 | 13.38 | 0.85 | 0.84 |
| LCO | 34.18 | 0.57 | 0.57 | 23.76 | 0.72 | 0.71 | 23.76 | 0.72 | 0.71 |
| LODO | 29.46 | 0.63 | 0.64 | 26.12 | 0.68 | 0.67 | 25.96 | 0.67 | 0.66 |
| LDO | 40.10 | 0.45 | 0.47 | 41.62 | 0.48 | 0.47 | 35.34 | 0.52 | 0.50 |

Table S12: The non-linear synthetic synergy model on the non-repeated dataset setup. Predictive under different L1 penalties ($\lambda$) and evaluation splits.

| **Split Type** | $\lambda = 0.1$ | | | $\lambda = 0.01$ | | | $\lambda = 0.001$ | | |
|---|---|---|---|---|---|---|---|---|---|
| | **MSE** | **PCC** | **SCC** | **MSE** | **PCC** | **SCC** | **MSE** | **PCC** | **SCC** |
| Non-repeated | 34.62 | 0.67 | 0.67 | 34.47 | 0.67 | 0.66 | 34.00 | 0.67 | 0.66 |

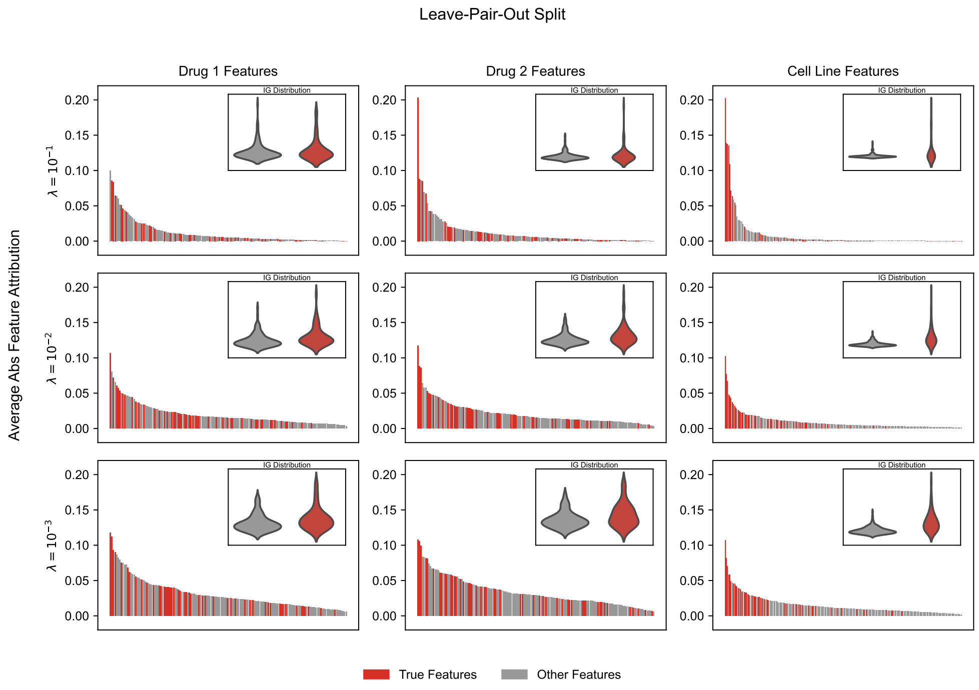


Figure S9: Integrated Gradients (IG) feature attribution for Leave-Pair-Out (LPO) split across L1 regularization values $\lambda = 10^{-1}, 10^{-2}, 10^{-3}$. Red bars represent informative features used during score generation; gray bars represent non-informative features. Despite strong predictive performance, attribution patterns show limited alignment with ground truth, indicating reliance on shortcut features rather than true informative signals.

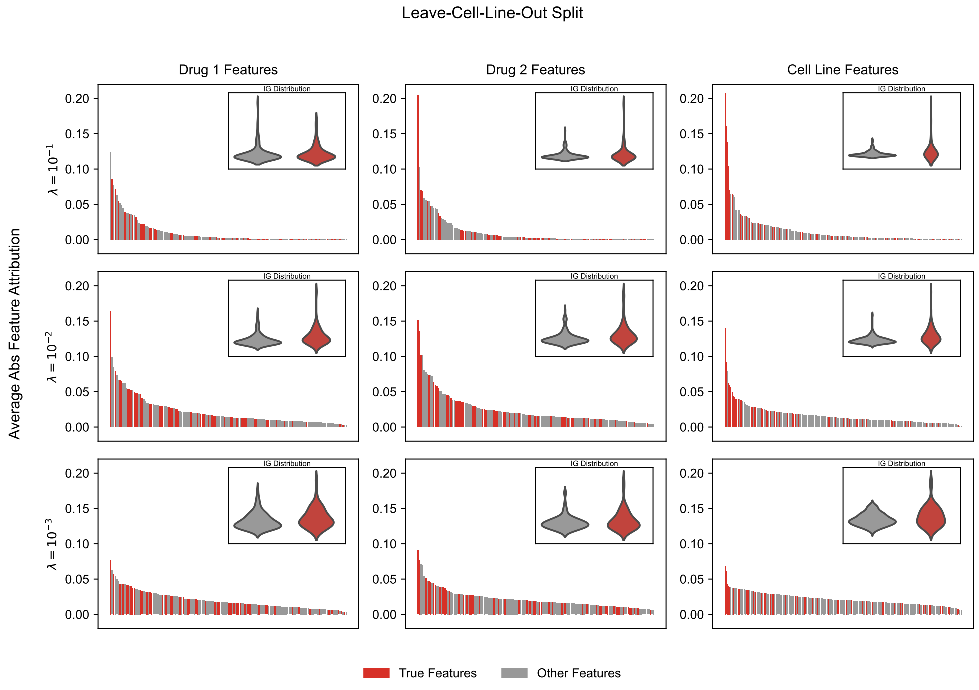


Figure S10: IG feature attribution for Leave-Cell-Out (LCO) split across L1 regularization values $\lambda = 10^{-1}, 10^{-2}, 10^{-3}$. Red bars represent informative features used during score generation; gray bars represent non-informative features. Similar to LODO, attribution does not consistently prioritize informative features, reinforcing evidence of shortcut learning under entity-based splits.

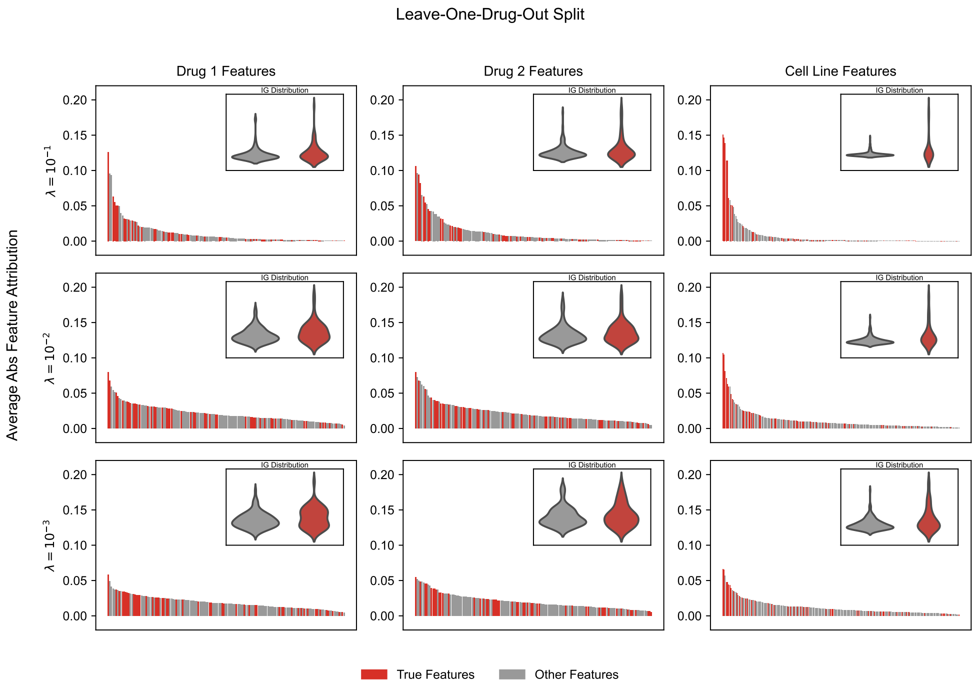


Figure S11: IG feature attribution for Leave-One-Drug-Out (LODO) split across L1 regularization values $\lambda = 10^{-1}, 10^{-2}, 10^{-3}$. Red bars represent informative features used during score generation; gray bars represent non-informative features. Models exhibit scattered attention across irrelevant features, suggesting poor feature-based generalization when encountering unseen drugs.

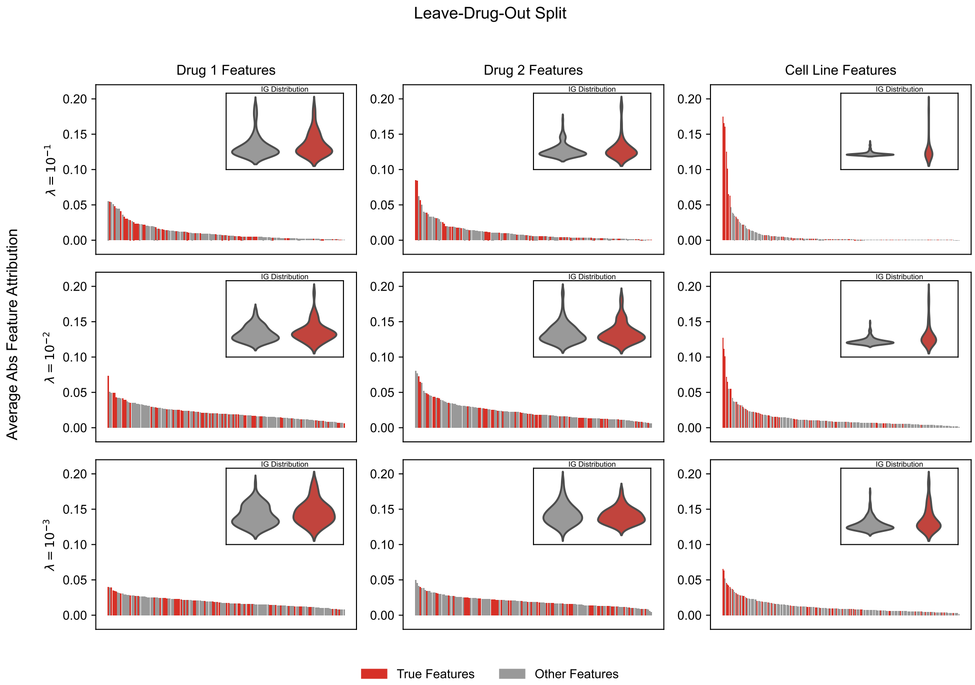


Figure S12: IG feature attribution for Leave-Drug-Out (LDO) split across L1 regularization values $\lambda = 10^{-1}, 10^{-2}, 10^{-3}$. Red bars represent informative features used during score generation; gray bars represent non-informative features. The model assigns importance broadly across non-informative features, showing minimal focus on the ground-truth informative set, suggesting reliance on dataset-specific co-occurrence cues and poor generalization to unseen drugs.

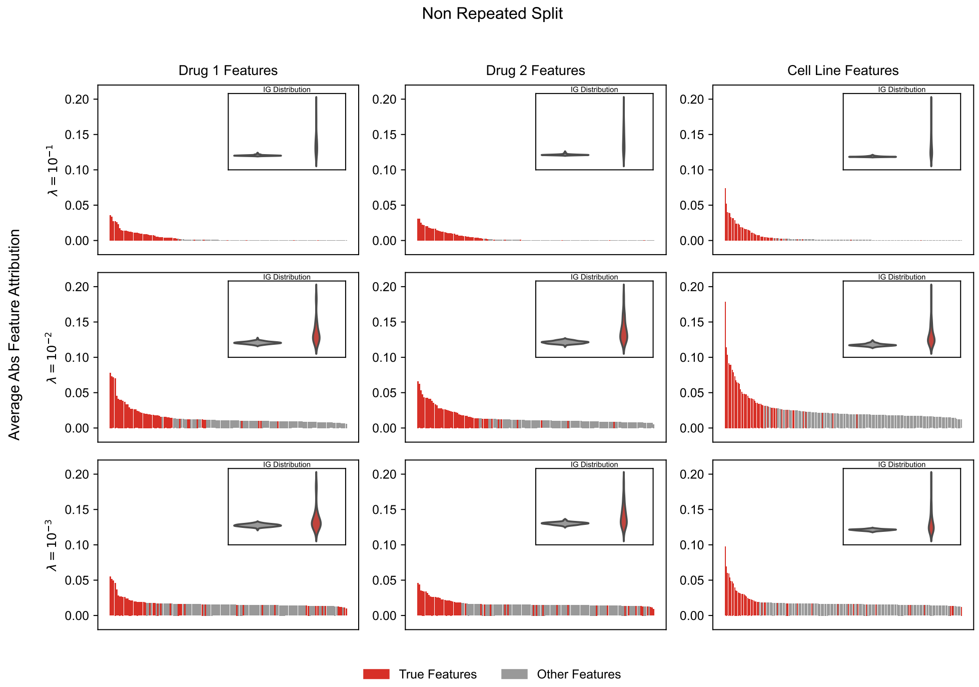


Figure S13: IG feature attribution for the non-repeated split across L1 regularization values $\lambda = 10^{-1}, 10^{-2}, 10^{-3}$. Red bars represent informative features used during score generation; gray bars represent non-informative features. Unlike repeated splits, models strongly prioritize informative features, as confirmed by a high Jaccard Index demonstrating that in the absence of shortcut opportunities, models learn true feature–outcome relationships.

### 9 Training Count-based Regression Trees

To understand entity pair count relations to the model error, we fit depth-2 regression trees across all applicable folds using Matchmaker results on DrugComb dataset. Shown in Figures S14, S15 and S16.

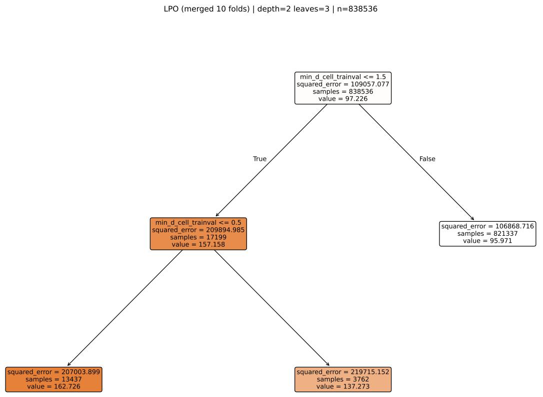


Figure S14: Depth-2 regression tree for the LPO split (10 folds merged). The tree uses the binned minimum train+val frequency of the two drug–cell pairs as input features. Very rare drug–cell pairs (min frequency is 0) show higher mean squared error, while the large majority of test examples fall into a leaf whose mean error is close to the global average.

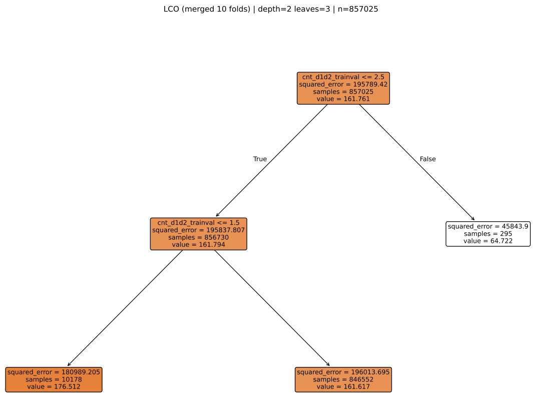


Figure S15: Depth-2 regression tree for the LCO split (10 folds merged). The tree uses only the binned train+val count of the drug pair as input. Pairs that are observed only a few times in train+val have higher errors on average, whereas most pairs with higher counts have mean errors similar to the global average.

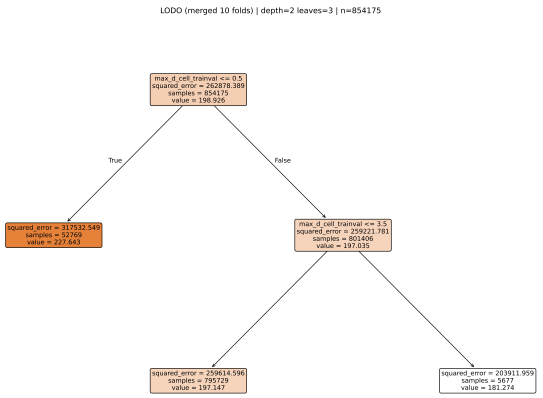


Figure S16: Depth-2 regression tree for the LODO split (10 folds merged). The tree is fitted using only the binned maximum train+val frequency of the two drug–cell pairs. Test examples with very low maximum drug–cell frequency tend to have higher errors, but for most examples the mean error remains close to the overall average across frequency bins.
